# Dominant Malignant Clones Leverage Lineage Restricted Epigenomic Programs to Drive Ependymoma Development

**DOI:** 10.1101/2024.08.12.607603

**Authors:** Alisha Kardian, Hua Sun, Siri Ippagunta, Nicholas Laboe, Hsiao-Chi Chen, Erik Emanus, Srinidhi Varadharajan, Tuyu Zheng, Blake Holcomb, Patrick Connelly, Jon P. Connelly, Michael Wang, Kimberley Lowe, Shondra M. Pruett-Miller, Kelsey C. Bertrand, Benjamin Deneen, Stephen C. Mack

**Affiliations:** Center of Excellence in Neuro-Oncology Sciences, St Jude Children’s Research Hospital, Memphis, TN, USA; Neurobiology and Brain Tumor Program, St Jude Children’s Research Hospital, Memphis, TN, USA; Department of Developmental Neurobiology, St Jude Children’s Research Hospital, Memphis, TN, USA; Cancer and Cell Biology Program, Baylor College of Medicine, Houston, TX, USA; Cell and Gene Therapy Program, Baylor College of Medicine, Houston, TX, USA; Center for Cancer Neuroscience, Baylor College of Medicine, Houston, TX, USA; Center for Advanced Genome Engineering, St Jude Children’s Research Hospital, Memphis, TN, USA; Department of Cell and Molecular Biology, St Jude Children’s Research Hospital, Memphis, TN, USA; Department of Oncology, St Jude Children’s Research Hospital, Memphis, TN, USA

## Abstract

ZFTA-RELA is the most recurrent genetic alteration seen in pediatric supratentorial ependymoma (EPN) and is sufficient to initiate tumors in mice. Despite ZFTA-RELA’s potent oncogenic potential, *ZFTA-RELA* gene fusions are observed exclusively in childhood EPN, with tumors located distinctly in the supratentorial region of the central nervous system (CNS). We hypothesized that specific chromatin modules accessible during brain development would render distinct cell lineage programs at direct risk of transformation by ZFTA-RELA. To this end, we performed combined single cell ATAC and RNA-seq analysis (scMultiome) of the developing mouse forebrain as compared to ZR-driven mouse and human EPN. We demonstrate that specific developmental lineage programs present in radial glial cells and regulated by Plagl family transcription factors are at risk of neoplastic transformation. Binding of this chromatin network by *ZFTA-RELA* or other PLAGL family motif targeting fusion proteins leads to persistent chromatin accessibility at oncogenic loci and oncogene expression. Cross-species analysis of mouse and human EPN reveals significant cell type heterogeneity mirroring incomplete neurogenic and gliogenic differentiation, with a small percentage of cycling intermediate progenitor-like cells that establish a putative tumor cell hierarchy. *In vivo* lineage tracing studies reveal single neoplastic clones that aggressively dominate tumor growth and establish the entire EPN cellular hierarchy. These findings unravel developmental epigenomic states critical for fusion oncoprotein driven transformation and elucidate how these states continue to shape tumor progression.

**HIGHLIGHTS:** 1. Specific chromatin modules accessible during brain development render distinct cell lineage programs at risk of transformation by pediatric fusion oncoproteins.

2. Cross-species single cell ATAC and RNA (scMultiome) of mouse and human ependymoma (EPN) reveals diverse patterns of lineage differentiation programs that restrain oncogenic transformation.

3. Early intermediate progenitor-like EPN cells establish a tumor cell hierarchy that mirrors neural differentiation programs.

4. ZFTA-RELA transformation is compatible with distinct developmental epigenetic states requiring precise ‘goldilocks’ levels of fusion oncoprotein expression.

5. Dominant tumor clones establish the entire EPN cellular hierarchy that reflects normal gliogenic and neurogenic differentiation programs.

## INTRODUCTION

Pediatric cancers are characterized by silent genomes, with fewer disease-driving mutations than their adult counterparts [1, 2]. Aberrant epigenomes play a predominant role in tumor initiation and development of many childhood cancers[3]. Fusion oncoproteins (FOs) are common drivers of pediatric cancers and often define their own disease subtypes, such as *EWSR1-FLI1* in Ewing sarcoma[4], *PAX3-FOXO1* in rhabdomyosarcoma[5], *ETV6-NTRK3* in infantile fibrosarcoma[6], and *ZFTA-RELA* in ependymoma[7]. While significant research has focused on molecular characterization of these FOs, the mechanisms of FO-driven transformation in the context of unique childhood developmental programs is incompletely understood. Specifically, how these FOs uniquely intersect and rewire developmental epigenetic states to transform cells remains unclear. To elucidate these mechanisms, we focused on ependymoma (EPN), the third most common pediatric brain tumor as a disease model. EPN are aggressive chemo-resistant pediatric brain tumors characterized by ‘quiet’ genomes, with very few recurrent genetic alterations, yet harbor aberrant tumor epigenomes [7, 8]. EPN that arise in the cortical brain (i.e. supratentorial, ST) are driven by a single gene fusion involving *ZFTA* and *RELA* (denoted ZR). *ZFTA* gene fusion events (including ZR) are observed near exclusively in EPN and occur specifically in the brain cortex, suggesting a direct relationship between ZR transformation and cortical progenitor populations [7]. EPNs are reported to arise from a type of radial glial cell (RGC) or gliogenic progenitor cell, present during embryonic brain development[9–11]. RGCs are multi-potent and can give rise to neurons and glia in two waves of differentiation during embryonic development[12–16]. While EPN cells express many RGC markers[9, 10, 17], the underlying molecular basis that places these cells at risk of transformation is not understood. In this study, we sought to understand the epigenomic landscapes of developmental cell lineages that render specific cell types vulnerable to transformation by FOs such as ZR. This was accomplished by dual single cell characterization of transcriptional and chromatin accessibility programs (termed scMultiome) of the developing mouse brain at embryonic day 14.5 (E14.5), a time of rapid RGC expansion, as well as integrated characterization across mouse and human EPN tumors. We applied *in vivo* barcoding technology to interrogate ZR-driven mouse EPN models and dissect the dynamics of malignant clonal diversity through integrated characterization of single cell transcriptional programs. Our investigation of the scMultiome landscapes of EPN across development and disease provide valuable insights into the cell lineage epigenomic programs that shape EPN initiation and progression. These findings may have broader ramifications upon our understanding of how pediatric fusion oncoproteins leverage distinct chromatin modules in distinct cell lineage programs to drive tumor development and cellular heterogeneity.

## RESULTS

### Lineage-specific chromatin modules at risk of transformation during embryonic development

ZFTA-RELA (ZR) functions as an oncogenic transcription factor (TF), activating neoplastic gene expression programs[17–19]. This finding led us to hypothesize that ZR remodels the developing epigenome and patterns of chromatin accessibility to activate oncogene expression. To test this hypothesis, we isolated putative EPN cells of origin, RGCs[7, 10], from Fabp7-eGFP mice and transduced cells with lentiviral vectors expressing ZR or an empty vector as a control (**Figure 1a**). ZR expression led to activation of *ZR* transcriptional targets and expression of a core 93 gene signature associated with ZR oncogenic activity (**Extended Data Figure 1a**)[17–19]. Unexpectedly, very minimal changes in patterns of chromatin accessibility were observed in ZR transformed RGCs profiled by ATAC-seq (**Figure1b**). These findings, instead, suggested that patterns of chromatin accessibility were already established in specific lineage programs and subsequently commissioned by ZR fusion oncoprotein binding.

**Figure 1.**
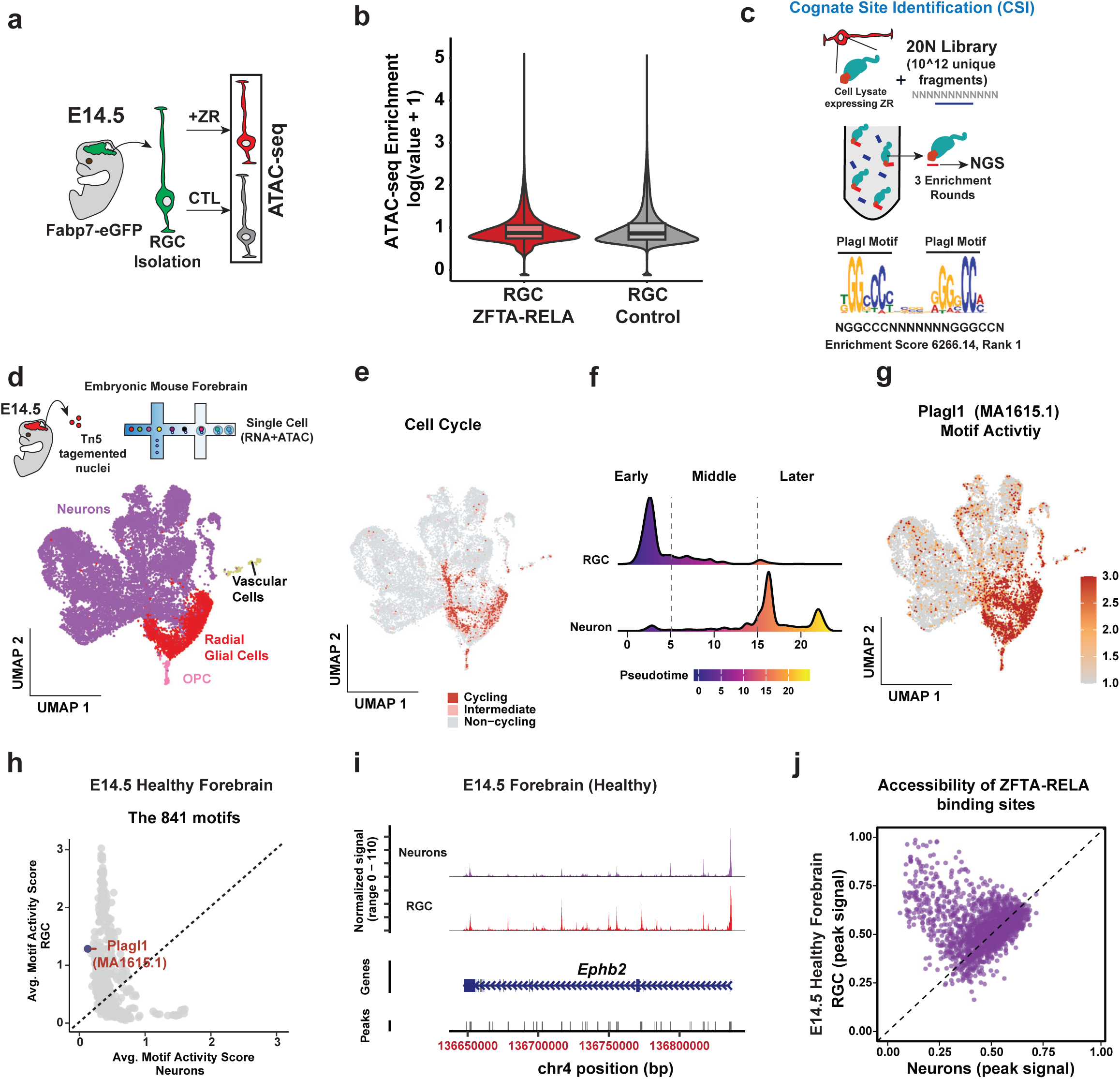
Fusion oncoproteins hijack lineage-restricted chromatin accessibility programs at risk of transformation during brain development. (**a**) Schematic illustrating the procedure for generating ATAC-Seq data. (**b**) Boxplots presenting global changes in chromatin accessibility patterns in control versus ZR-transformed RGCs. (**c**) A schematic illustrating the cognate site identification (CSI) method for analyzing the binding affinity of ZR for *PLAG* family motifs. (**d**) Schematic depicting the methodology for generating scMultiome data from E14.5 normal mouse forebrain. The UMAP plot illustrates the distribution of cell types observed following analysis of the scMultiome data. (**e**) Patterns of cell cycling and non-cycling states in the E14.5 normal mouse forebrain. (**f**) Estimating molecular pseudo-time for snRNA data and plotting cell density at different time intervals against pseudo-time for each cell. Time intervals: 0-5 for “Early”, 5-15 for “Middle”, and >15 for “Later”. (**g**) UMAP reveals Plagl1 motif activity patterns in E14.5 normal mouse forebrain via scMultiome data. (**h**) Comparison between RGC and neurons across 841 global motif activities, with the Plagl1 motif highlighted. Each dot represents an average motif activity. (**i**) Pseudo-batch accessibility trace from snATAC, where signals from RGC and neurons within the group have been averaged together to visualize DNA accessibility in the Ephb2 gene region. (**j**) The chromatin accessibility ratio of ZR binding sites between RGCs and neurons. Each point represents a ZR binding peak, and the x/y axis represents the percentage of cells with an open ZR binding region.

While previous CUT&RUN experiments suggest PLAGL1/2 binding motifs are shared by ZR, the exact binding sequence of the fusion remains unknown[17]. To understand the DNA sequence binding preferences of ZR directly, we adapted the cognate site identification (CSI) assay[20, 21] for use with ZR-expressing protein lysates. This involved immunoprecipitation of HA-tagged ZR-expressing cell lysates with a library of 1 trillion unique combinations of double-stranded DNA fragments across a stretch of 20 base pairs (bps). Enrichment of DNA fragments was performed followed by PCR amplification and Illumina short-read sequencing. Amplified fragments were then used in subsequent rounds of immunoprecipitation for a total of 3 rounds. CSI profiling of ZR revealed a very strong binding preference for Plagl family TF motifs engaged in a dimerized fashion, and enrichment of a core GGGCC consensus binding sequence (**Figure 1c, Extended Data Figure 1d**). We therefore asked to what extent are Plagl family TF motifs accessible during embryonic brain development, and in what developmental cell types.

To this end, we micro-dissected forebrains from mouse embryos day E14.5, a timepoint during active RGC expansion[22, 23]. Tissue was dissociated and subjected to single cell ATAC- and RNA-seq from the same nuclei, using the scMultiome assay developed by 10x Genomics (**Figure 1d**). Cell types were annotated using a previously established developmental mouse brain atlas[23] (**Extended Data Figure 1b,e**; **Table 2S**), revealing RGCs, neurons, OPCs, immune cells, and vascular cells, along with their transcriptomic and epigenomic signatures (**Figure 1d; Table S3**). As expected, RGCs were actively proliferating, giving rise to post-mitotic neurons (**Figure 1e-f**) during neurogenesis. Critically, we found that the Plagl family TF motifs were accessible preferentially in the RGC lineage, and chromatin accessibility was reduced upon neuronal differentiation (**Figure 1g-h; Extended Data Figure 1g-h**). An examination of ZR specific DNA binding sites revealed cell type specific accessibility in RGCs versus neurons, including known ependymoma oncogenes such as *EphB2* and *Notch1* (**Figure 1i-j; Extended Data Figure 1c,i**). These findings pinpoint chromatin signatures at direct risk of transformation in distinct cell lineages present during brain development as an underlying basis for EPN initiation.

### Developmental programs establish tumor cell heterogeneity and restrain EPN cell proliferation

To study the intersection of developmental and ZR-driven oncogenic programs in EPN, we leveraged a natively forming ZR EPN mouse model established by *in utero* electroporation (IUE) and characterized tumors using scMultiome profiling[24–26]. This involved stable integration of DNA transgenes expressing ZR using the PiggyBac (PB) transposon system at E16.5; targeting RGCs with a PB transposase vector driven by a *Glast* promoter[17] (**Figure 2a**). Using IUE, we generated a series of ZR driven EPN and glioblastoma (GBM) mouse tumors for comparative analyses (**Figure 2b-c**). EPN tumors were established by expression of ZR, and GBM by simultaneous CRISPR-Cas9 mediated knockout of *Nf1;Pten;Tp53* at E16.5 (**Figure 2b-c**, denoted 3xCR)[27, 28].

**Figure 2.**
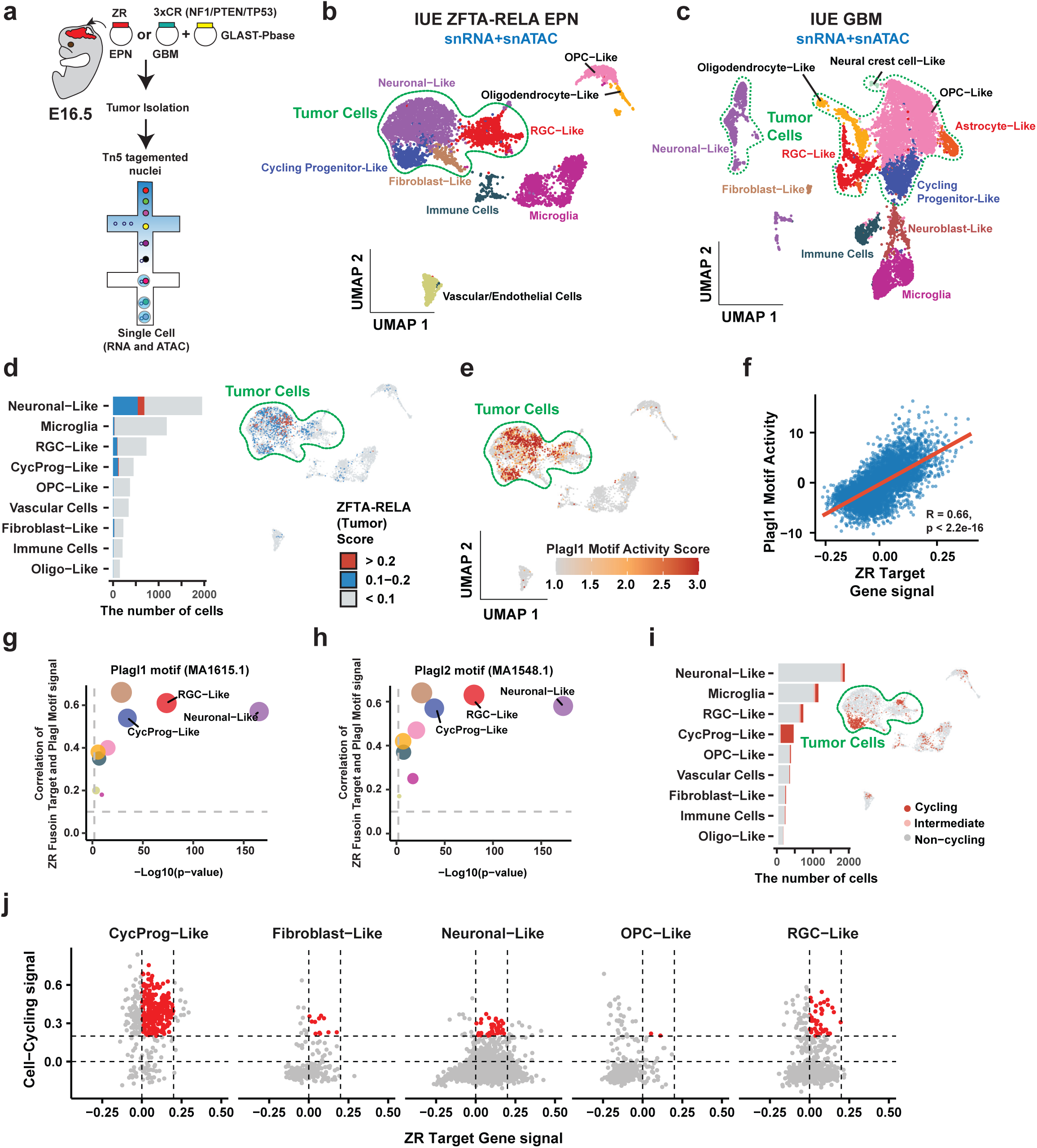
ZFTA-RELA fusion signature is enriched in neuron-like cells in mouse IUE. (**a**) Workflow for mouse IUE tumor generation and preparation of scMultiome libraries and data. (**b**) UMAP plot of three integrated ZR tumors generated by IUE. Tumor cells are identified by predicting copy numbers from snATAC data. (**c**) UMAP plot of glioblastoma tumor generated by IUE. (**d**) Bar graph for ZR signal in each cell type and UMAP for the ZR signal in each individual cell. ZR activity fraction for each cell was estimated based on the gene expression of a set of 93 genes associated with ZR activity. >0.2 indicates a strong ZR signal, 0.1-0.2 indicates a moderate ZR signal, and <0.1 indicates no detectable ZR signal. (**e**) Plagl1 motif activity per cell in IUE ZR tumors. (**f**) Correlation between Plagl1 motif activity and ZR signatures across all cells. The x-axis is the ZR signal score, and the y-axis is the Plagl1 motif activity score. (**g**) Correlation (y axis) and significance (x axis) of Plagl1 motif activity and ZR signatures across various cell types. (**h**) Correlation (y axis) and significance (x axis) of Plagl2 motif activity with ZR signatures across various cell types. (**i**) Bar graph for the cell cycling signal in each cell type and UMAP for the cell cycling signal in each individual cell. (**j**) Comparing cell cycling signal score (y axis) to ZR signature score (x axis) across the various cell types.

The developmental mouse brain atlas that was used to identify cell types in normal development (**Figure 1**) was also applied to EPN and GBM mouse tumors. We observed diverse patterns of malignant and non-malignant cell types between ZR EPN and 3xCr GBM, and the absence of a complete differentiation ‘block’ that would be consistent with accumulation of specific cell types transformed during embryonic brain development (**Figure 2b-c; Extended Data Figure 2a-g; Table S4**)[29]. Instead, mouse ZR EPN tumors displayed a diversity of malignant cells with signatures like RGCs, cycling neural intermediate progenitor cells (IPCs), fibroblast cells, and neuronal cells (denoted neuronal-like) (**Figure 2b**). Interestingly, very few cells along the oligodendrocyte progenitor cell (OPC) to oligodendrocyte lineage were associated with neoplastic transformation, as compared to mouse GBM tumors that demonstrated predominant cellular expansion of the OPC lineage (**Figure 2c**)[30, 31].

Application of the ZR 93 gene target signature (**Figure 1b**), could also robustly identify malignant cells that corroborated tumor cell annotation using copy number variants (**Figure 2d; Extended Data Figure 2a-c**). This analysis revealed wide-spread ZR target activity across all malignant cell states detected within RGC-like, cycling progenitor-like, and neuronal-like cells. In sharp contrast to normal mouse forebrain development (**Figure 1g**), Plagl1 motif activity (a metric based on motif occurrence and gene expression, please see Methods) was wide-spread across all malignant cell types and highly correlated with ZR target activity (**Figure 2e-f**). This contrast was further highlighted by the correlation between ZR and Plagl1 target activity observed preferentially in neuronal-like EPN cells (**Figure 2g-h; Extended Data Figure 2h-m**).

Finally, we quantified the cell cycling state across different developmental cells identified in ZR driven EPN mouse tumors (**Figure 2i-j**). This revealed that only a sub-population of cycling progenitor cells were actively dividing, with other cell types being mostly non-proliferative. Interestingly, when compared against ZR activity, only cycling progenitors with low to moderate ZR activity exhibited the highest cell proliferation signal (**Figure 2j**). This was in contrast with other developmental cell types (i.e. neuronal-like) with much fewer cells in a proliferative state despite the presence of ZR oncogenic activity. Together, these data demonstrate that the transcriptional diversity of ZR ependymoma mirrors normal (albeit incomplete) developmental programs, and that differentiated cell types are resistant to ZR-induced proliferation because of their more restricted epigenomic states.

### Convergent activation of developmental PLAGL TF motifs by pediatric fusion oncoproteins

We sought to corroborate our findings in murine EPN by studying human EPN samples. Primary tumors from 19 patient samples, including 9 supratentorial ependymomas (ST-EPN) and 10 posterior fossa ependymomas (PF-EPN) were profiled by scMultiome analysis (**Figure 3a-b**). Upon cell type annotation, non-tumor cells included endothelial cells, immune cells (microglia and other immune cells), and oligodendrocytes. We defined malignant and non-malignant cells based on the copy number alterations (CNAs) detected in each cell (**Figure 3c; Extended Data Figure 3a-b,** See Methods). Using 241 gene expression markers established by *Pajtler* et al. 2015[32], tSNE profiling of scRNA data segregated PF ependymomas, and resolved two groups of ST ependymomas (**Figure 3d-f**). DNA methylation array profiling showed that these two groups of ST tumors harbored *ZFTA-RELA* or *EWSR1-PLAGL1* (denoted EWP) gene fusions, both with intact DNA-binding domains and predicted to bind Plagl family DNA motifs (**Figure 3e**). There were significant differences in the gene expression signatures of PF EPN, and those driven by ZR or EWP fusions (n=307 genes, adjusted p-value < 0.001). Notably, expression of at least 20 TFs were significantly different between the three groups, such as *PAX3, MECOM* and *AR* in PF ependymoma, *LHX2* and *RELA* in ZR EPN, and *PAX2* and *PAX7* in EWP tumors (**Figure 3f; Table S5**). Consistent with murine ZR EPN, we found that ZR fusion activity per cell could be predicted across species using a common 93 gene signature and used to robustly stratify ZR driven EPN tumors (**Figure 3g)**.

**Figure 3.**
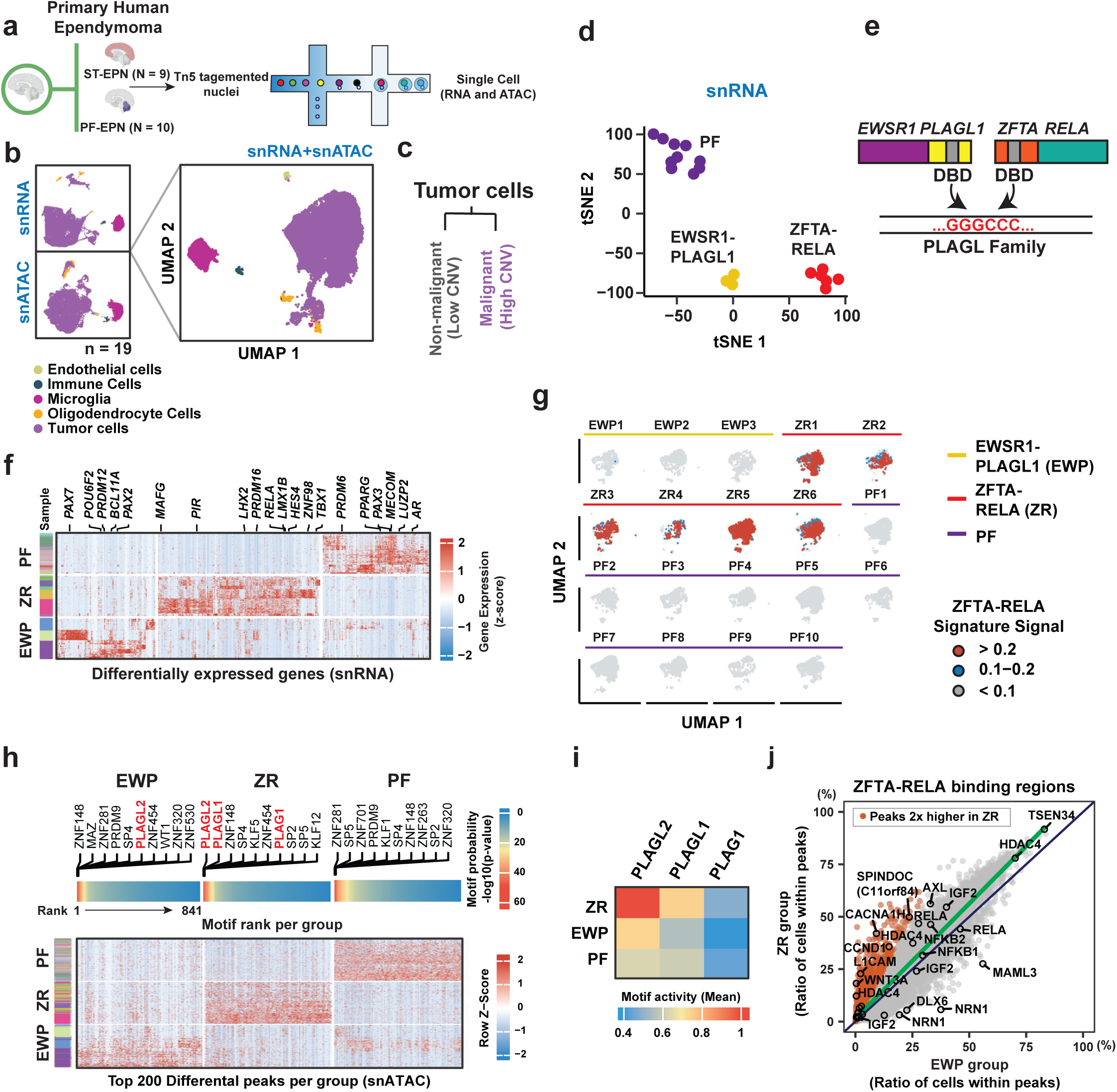
PLAGL family motifs are highly enriched in human ZFTA-RELA EPN compared with other EPN subgroups. (**a**) Workflow for the generation of supratentorial (ST-EPN) and posterior fossa (PF-EPN) human scMultiome sequencing data. (**b**) scMultiome data from 19 patients, along with initial cell type annotation. UMAPs were generated from snRNA-, snATAC-, and scMultiome-integrated data. (**c**) Tumor cells can be classified into non-malignant and malignant types according to their degree of copy number variation. (**d**) Analysis of malignant tumor cells based on gene expression markers shows our EPNs can be classified into three molecular subgroups (PF, EWSR1-PLAGL1, and ZFTA-RELA). Specific subgroup characteristics were further validated through DNA methylation analysis. (**e**) Diagram illustrating the convergence of different EPN fusions on PLAGL family motifs via their DNA binding domains, suggesting a shared mechanism of action. (**f**) Heatmap illustrating genes with significant differences in expression among the three EPN subgroups (including EWP-, ZR-, and PF-group) within malignant cells. Transcription factors demonstrating significant differences are highlighted. (**g**) ZR fusion signals were assessed in malignant cells across each sample via expression of the 93 gene signature used previously. (**h**) Heatmap displaying the top 200 differential snATAC peaks for each group. The upper heatmap illustrates enriched motifs ranked from highest to lowest among these top 200 peaks, with the top 10 gene names of these motifs provided. The PLAG family motifs are highlighted in red. (**i**) Average activity of PLAG family motifs across EPN subgroups. (**j**) The proportion of cells in which the ZR binding region is open between the EWP and ZR groups. Orange binding regions are significantly higher in the ZR group.

Next, we determined whether ZR and EWP tumors converge upon shared chromatin accessibility programs enriched in developmental PLAGL family motifs. As expected, the PLAGL family (*PLAG1, PLAGL1, PLAGL2*) motifs were highly enriched in the ZR group among the top 200 differential scATAC peaks (**Figure 3h; Table S5**). Of note, PLAGL2 motifs were also highly enriched amongst the top DNA motifs in EWP tumors, as compared to PF EPN. Comparing the PLAGL family motif activity across tumors, we found the highest levels in ZR ependymoma, followed by increased activity in EWP tumors, and the lowest levels in PF EPN (**Figure 3i)**. This data suggests shared utilization of PLAGL family motifs, and distinct oncogenic targets between ZR and EWP tumors (**Extended Data Figure 3c-e; Table S5**). This concept was supported by direct comparison of scATAC data between ZR and EWP tumors with a focus on known ZR binding sites **(Fig 3j)**. While many genomic regions were shared between ZR and EWP tumors suggesting convergent oncogenic mechanisms, highly expressed signature genes of ZR ependymoma such as *L1CAM*, *CCND1*, *WNT3A* were accessible specifically in ZR EPN. These findings underscore the utilization and diversity of PLAGL TF family motifs that are leveraged by pediatric fusion oncoproteins to drive brain tumor development.

### Human EPN cellular diversity is consistent with murine EPN and mirrors specific lineage programs

We next characterized the cell types present in ST and PF ependymoma along with their patterns of chromatin accessibility. Since ST-EPN occurs in the supratentorial region and PF-EPN occurs in the posterior fossa (including brainstem and cerebellum), we annotated malignant cells with different human cell atlases of the developing human brain[33] (**Table S2; Extended Data Figure 4a-b,k**). Of the total number of malignant cells in ST-EPN, 69% and 31% of cells belonged to ZR group and EWP group respectively. Consistent with murine EPN, scMultiome analysis revealed malignant cells classified as neuronal-like, RGC-like, astrocyte-like, OPC-like, and nIPC-like. Similarly, PF EPN was composed of 7 different cell types, such as neuronal-like, RGC-like, astrocyte-like, OPC-like, nIPC-like, and the additional presence of choroid plex (CP)-like and ependymal-like cells (**Figure 4a-d; Extended Data Figure 4c-d; Table S6**). Given the over-representation of PLAGL TF family motifs in ST EPN, we next sought to investigate the enrichment of these DNA motifs within EPN cell populations. In ZR and EWP driven tumors, we found that PLAGL2 motif activity was most elevated in neuronal-like cells, with accessibility also in nIPC-like and RGC-like cells (**Figure 4e**).

**Figure 4.**
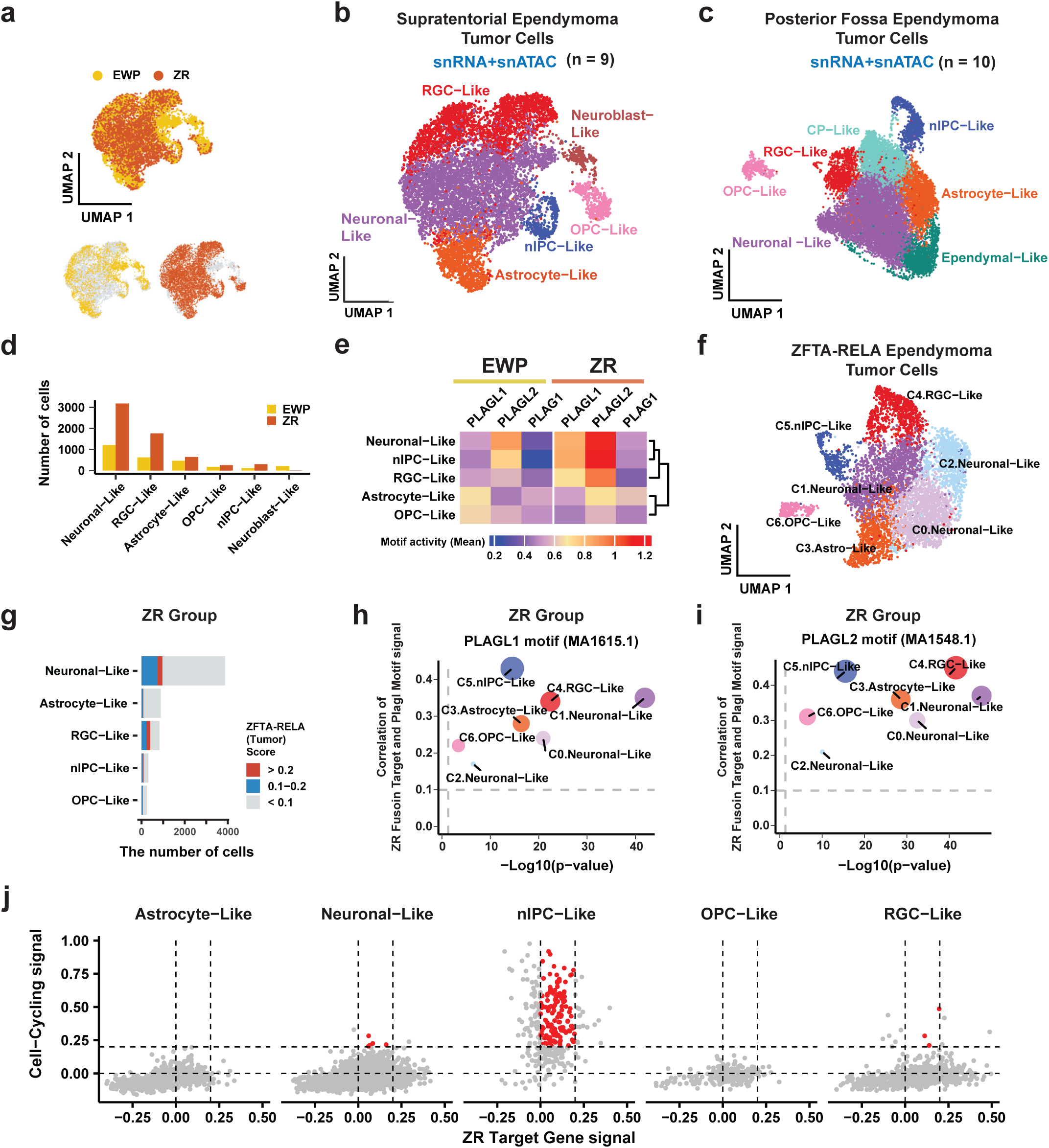
ZFTA-RELA and PLAGL family motifs are highly active in neuron-like cells in patient samples. (**a**) Integrated scMultiome UMAP for ST-EPN, including EWP and ZR groups. (**b**) Cell type annotation for integrated ST-EPN multiome data. (**c**) Cell type annotation for integrated PF-EPN scMultiome data. (**d**) The number of malignant cells in each cell type, compared between EWP and ZR groups. (**e**) Average activity of PLAG family motifs across different cell types within the EWP and ZR groups. (**f**) Distribution of cell types in the integrated ZR group scMultiome dataset. ‘C’ is an abbreviation for cluster. (**g**) Bar graph for ZR signal in each cell type and UMAP for the ZR signal in each individual cell. The ZR activity fraction for each cell was estimated based on the gene expression of a set of 93 genes associated with ZR activity. “>0.2” indicates a strong ZR signal, “0.1-0.2” indicates a moderate ZR signal, and “<0.1” indicates no detectable ZR signal. (**h**) Correlation (y axis) and significance (x axis) of Plagl1 motif activity and ZR signatures across various cell types. (**i**) Correlation (y axis) and significance (x axis) of Plagl2 motif activity and ZR signatures across various cell types. (**j**) Comparing cell cycling signal score (y axis) to ZR signature score (x axis) across the various cell types.

To further delineate ZR EPN and validate our mouse model, we separated the ZR group and re-clustered and re-annotated the data (**Figure 4f; Extended Data Figure 4e-i; Table S6**). Among ZR malignant cells, ZR signatures were highly detected in neuronal-like and RGC-like cells compared with other cell types (**Figure 4g**). ZR signatures were also positively correlated with PLAGL family motif activity (**Figure 4h** and **4i**). Like murine ZR EPN, positive correlation between ZR signature and PLAGL family motif activity were detected largely in neuronal-like cells. Compared to normal RGC development, this suggests that ZR expression maintains chromatin accessibility of PLAGL family motifs that are normally repressed during neuronal differentiation. Finally, we predicted cycling signatures and compared them with ZR signatures in different cell types (**Figure 4j; Extended Data Figure 4j**). Among them, only nIPC-like cells with low-moderate ZR activity exhibited the highest cell proliferation signal, thus highly consistent with ZR mouse EPN (**Figure 4j and Figure 2j**).

### Highly dominant tumor clones maintain EPN transcriptional heterogeneity in murine EPN

To study how developmental lineage programs intersect with tumor heterogeneity, we leveraged an *in vivo* barcoding system called Tracker-seq to label cells and read out barcodes using scRNA sequencing[34]. Tracker-seq was used to barcode tumor cells arising in a natively forming IUE mouse model of ZR EPN (**Figure 5a; Table S7**). Here, we examined two different timepoints, a postnatal day 4 (P4) sample, versus fully formed late-stage tumors (∼P70) (**Figure 5b, Extended Data Figure 5**). Significant tumor clonal diversity was observed in the early-stage neoplastic lesion with 4- to 6-fold greater numbers of lineage barcodes detected than endpoint tumors. In contrast, we detected the emergence of dominant largely single tumor cell clones in late-stage tumors, with the dominant lineage barcode detected in 89∼99% of the cells (**Figure 5c-e**). Critically, cell type annotation revealed that dominant tumor cell clones could establish the entire developmental and transcriptional diversity of EPNs seen in the mouse and human disease (**Figure 5f, 5i; Table S7**). Within these dominant tumor cell populations, the ZR signature was highly enriched in neuronal-like cells and cycling progenitor-like cells (**Figure 5g, 5j**). Consistent with scMultiome profiling of mouse and human ZR EPN, the cycling signature was highly active in cycling progenitor cells (**Figure 5h, 5k**). These findings reveal dominant tumor clones in EPN that drive tumor formation and can establish the entire cellular diversity of the disease.

**Figure 5.**
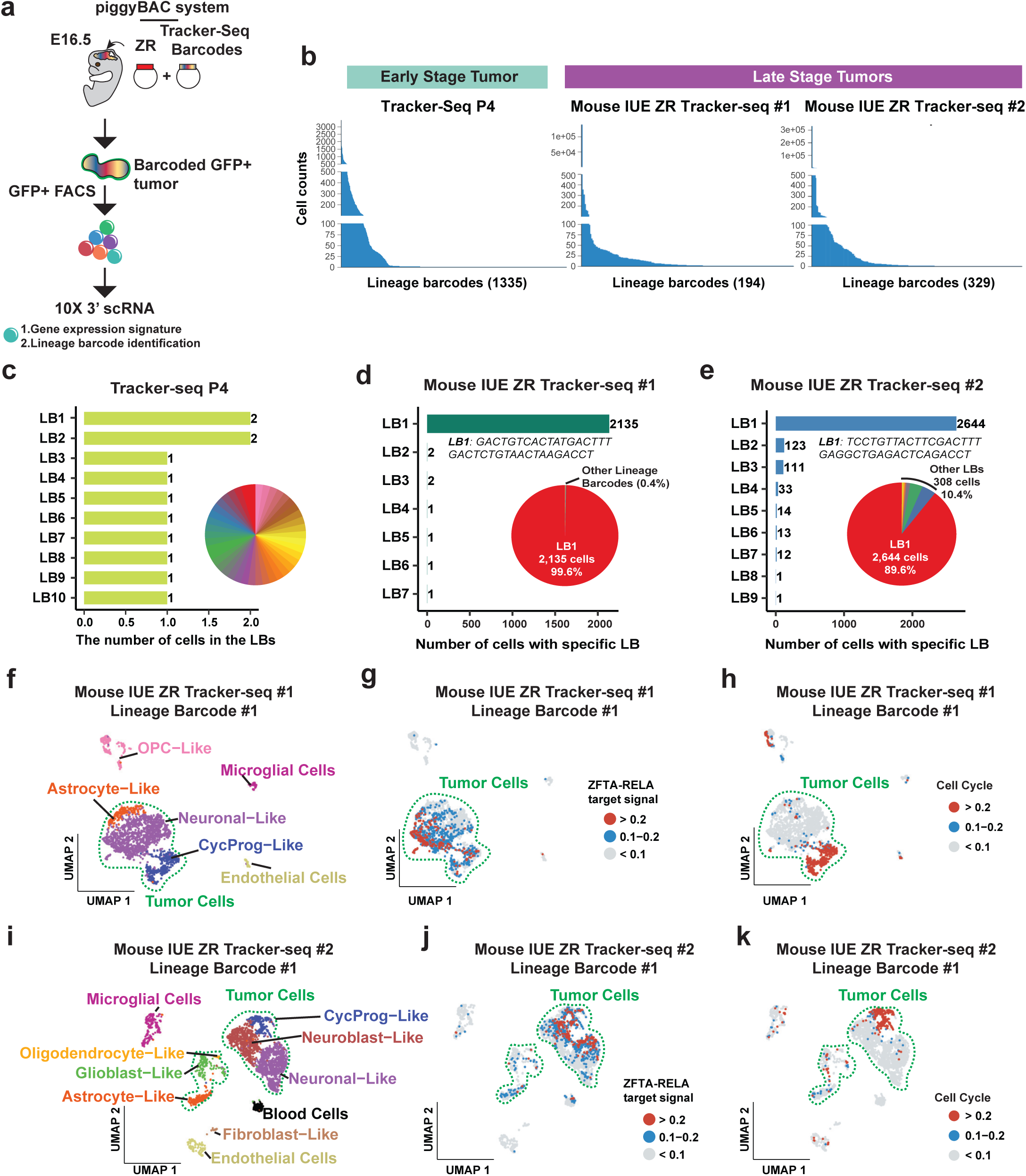
Single dominant ZFTA-RELA tumor clone can form independent lineage trajectory. (**a**) Schematic illustrating the procedure for generating Tracker-Seq data. (**b**) Count of unique lineage barcodes in the raw Tracker-Seq data and the number of cells associated with each lineage barcode. (**c-e**) Top 10 lineage barcodes ranked by the number of cells per barcode. ‘LB’ is an abbreviation for lineage barcode. This represents a filtered lineage barcode dataset, where cells with more than one barcode are removed. (**f** and **i**) Cell type distribution in the Tracker-Seq data for 2 biological replicates. (**g** and **j**) The ZR activity fraction for each cell was estimated based on the gene expression of a set of 93 genes associated with ZR activity. “>0.2” indicates a strong ZR signal, “0.1-0.2” indicates a moderate ZR signal, and “<0.1” indicates no detectable ZR signal. (**h** and **k**) Cell cycling activity fraction for each cell was estimated based on the gene expression. >0.2 indicates a strong signal, 0.1-0.2 indicates a moderate signal, and <0.1 indicates no detectable signal.

### Early progenitor-like cells govern EPN cell hierarchy and give rise to infrequently dividing cell progeny

To understand malignant cell lineage trajectories and which cell types develop into early and late stages of malignancy, we performed pseudotime analysis for each malignant cell type and constructed lineage models. pseudotime analysis of dominant Tracker-seq clones suggested a model in which cycling progenitor-like cells give rise to astrocyte-like cells or neuronal-like cells to mirror normal (albeit incomplete) neural differentiation programs. (**Figure 6a, 6d, 6g; Table S8**). Based on the Pseudotime evolutionary inference results, we generated a lineage trajectory model, centered upon cycling progenitor cells giving rise to at least two distinct cell lineage programs labeled ‘gliogenic’ versus ‘neurogenic’ (**Figure 6b, 6e; Table S8**). Analysis of cell cycle programs within this trajectory analysis supported a model of cycling progenitor cells giving rise to either slowly dividing or post-mitotic cells within the tumor microenvironment (**Figure 6c, 6f**). To confirm whether the lineage trajectory model derived from mouse IUE data could extend to patient tumors, we used human ZR group malignant cells and performed the same analysis. Pseudotime analysis demonstrated very similar patterns of nIPC-like cells arising early, and subsequently giving rise to gliogenic and neurogenic differentiation programs (**Figure 6g; Table S8**). The lineage trajectory model developed in murine ZR EPN applied to the human disease, revealing a similar pattern of cycling progenitor cells projected to differentiate into post-mitotic lineage programs (**Figure 6h-i**). Together our data support a model in which EPN cellular heterogeneity is established by dominant progenitor-like cells that give rise to post-mitotic or infrequently dividing glial- or neuronal-like cells.

**Figure 6.**
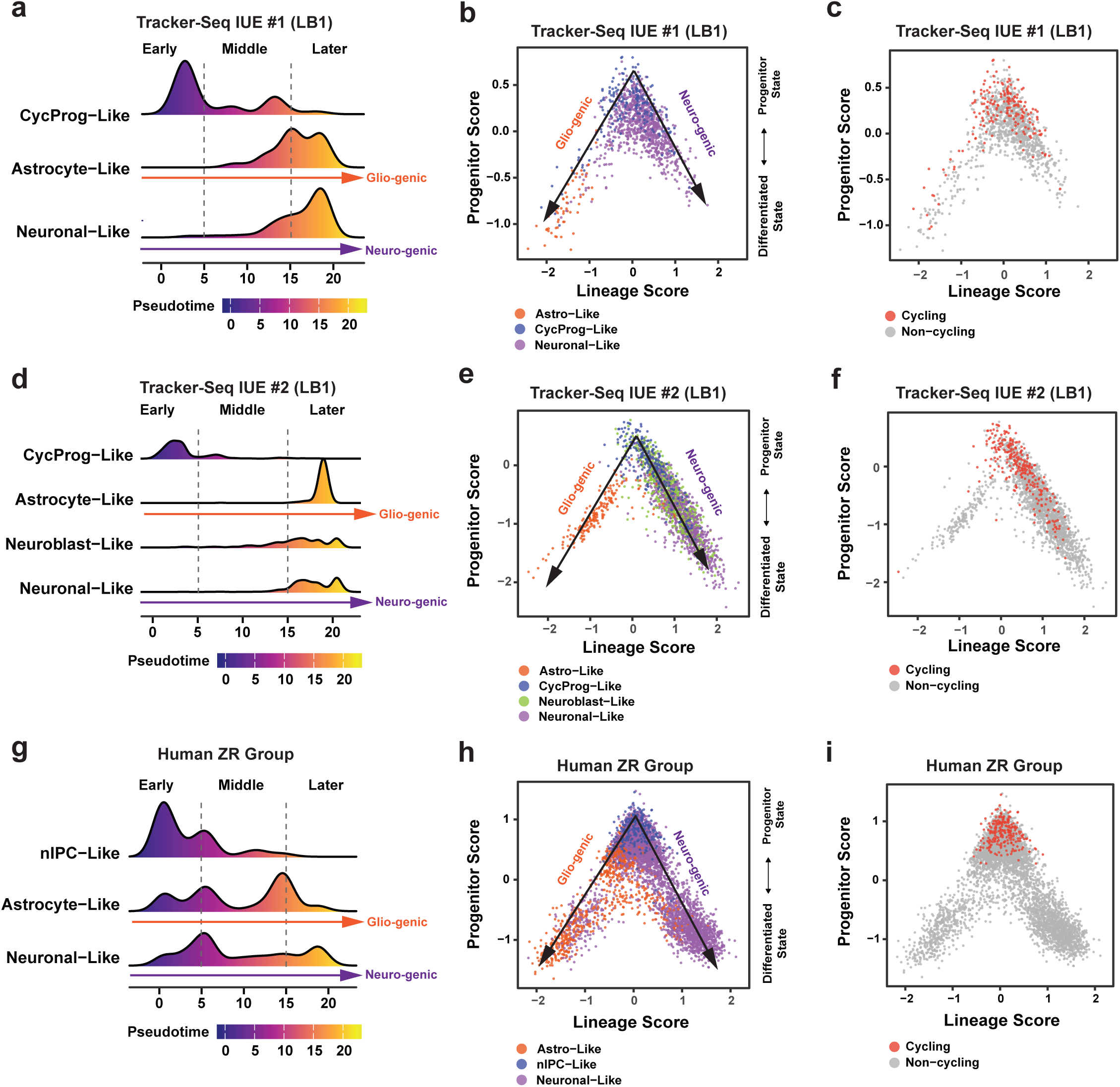
ZFTA-RELA tumor cycling cells are formed at an early stage and, like cancer stem cells, can differentiate into different cell types. (**a, d, g**) Cell density at different time intervals against molecular pseudo-time for each cell in Tracker-Seq and human datasets. Pseudo-time intervals: 0-5 for “Early”, 5-15 for “Middle”, and >15 for “Later”. (**b, e, h**) Triangular diagram showing malignant cell lineage and differentiation status. (**c, f, i**) Same triangular diagram showing malignant cell lineage and differentiation status, but with red dots indicating cells within the cell cycle.

## DISCUSSION

Pediatric brain cancers are often considered a disease of development[35], characterized by neoplastic transformation of early developmental cells. In ependymoma (EPN), multipotent RGCs have been hypothesized to be the candidate cells of origin[9, 10]. However, the underlying mechanistic basis for this connection has remained unclear. scMultiome characterization of the developing mouse brain reveals a specific chromatin accessibility program enriched in PLAGL family DNA motifs that is at direct risk of cellular transformation in developing RGCs. *ZFTA-RELA* gene fusions, which are observed exclusively in EPN, are not capable of remodeling chromatin accessibility, but instead engage developmental PLAGL family motifs that are already open in development at most oncogenic loci (i.e. *Ccnd1* and *Notch1*). PLAGL family genes play roles in corticogenesis and patterning, proliferation, and development[36–38] and are expressed in other cancer types[39–41] Considering the role of PLAGL1/2 in development and disease, the ability of ZR to keep these PLAGL family target genes open beyond embryonic stages may be an important mechanism in ST-EPN development and maintenance. As a result, pro-proliferation genes that govern RGC progenitor expansion are kept active, which normally become repressed as gliogenesis and neurogenesis progress. These findings uncover distinct epigenomic states within developing cell populations that establish direct risk of transformation by highly specific DNA motif targeting fusion oncoproteins, such as ZFTA-RELA.

Our findings may broadly implicate PLAGL family DNA motifs as direct targets in brain tumor transformation. Supporting this concept, we found shared chromatin accessibility programs that converged upon PLAGL family motifs when comparing ZR EPN against EWP ependymoma-like brain tumors. Interestingly, and despite similar PLAGL motif enrichment, there were significant differences in gene expression programs that distinguished ZR from EWP tumors, suggesting influence from developing cells of origin, or different target binding preferences between the two fusion oncoproteins. Beyond ZR and EWP, *PLAG1* is amplified and over-expressed in subsets of central nervous system (CNS) embryonal tumors and is one of the top up-regulated genes positively correlated with oncohistone histone H3 K27M diffuse midline glioma[39, 41, 42]. These observations lead us to hypothesize that the PLAGL transcription factor cistrome, and its targeting by specific oncoproteins, have broader significance in the development of a wide variety of pediatric brain tumors.

Our study evaluated the intersection of oncogenic and developmental programs from persistent ZR expression during brain development. scMultiome analysis of the ZR driven IUE model revealed diverse cell types with distinct transcriptional and chromatin accessibility programs. These findings were validated across species by scMultiome profiling of mouse and human ZR EPN. While we anticipated a significant ‘block’ in development at the RGC state, we were surprised to observe continued (albeit incomplete) differentiation through the astrocytic and neuronal lineages, with most differentiated cells in EPN classified as infrequently or non-cycling. Despite ZR fusion activity across malignant cell types, cell cycle signatures were seen prominently in the progenitor cell state, and rarely in differentiated populations such as neuronal-like and astrocyte-like cells. In normal brain development, RGCs often asymmetrically differentiate to intermediate progenitor cells (IPCs), which then terminally differentiate to neurons[43–45]. This could suggest a mechanism whereby a minority of transformed ZR driven cells accumulate in the IPC state, with most cells continuing to differentiate along the neuronal or glial cell lineage. Interestingly, the fact that IPCs are committed neural precursor cells could explain the high number or neuronal-like cells seen in these EPN tumors, compared to other tumors such as GBM, which appears to develop along the OPC lineage. These findings reveal the specific cellular transcriptional and epigenomic states compatible with ZR activity that drive EPN development. Interestingly, cell proliferation was most correlated with low- to modest-levels of ZR activity, suggesting like other pediatric FOs (i.e. EWS-FLI1), a ‘goldilocks’ principle that governs oncoprotein expression and neoplastic transformation potential [46, 47]. Together, these data have potentially important therapeutic applications predicated upon inducing cellular differentiation of progenitor-like tumor cells to adopt a more differentiated state that is post- or slowly-mitotic.

*In vivo* lineage tracing experiments of murine ZR EPN demonstrated the emergence of dominant (often single) clones, and the capability of such clones to establish the entire cellular diversity of ZR-driven EPN. Examples of clonal dominance in brain tumors have been observed, but often in adult tumors (such as glioblastoma) where clonal competition relies on successive mutations that are not often seen in pediatric brain tumor patients[48, 49]. Our findings suggest that upon expression of ZR, cellular diversity is established through existing developmental programs associated with distinct transcriptional and chromatin states. These findings have potential clinical ramifications for EPN as conventional therapies, such as surgery and radiotherapy, may miss single aggressive tumor clones capable of driving tumor recurrence.

Finally, based upon mouse and human ZR EPN, we propose a model in which a small proportion of cycling progenitor cells contribute to EPN initiation, and give rise to gliogenic and neurogenic lineage programs, and differentiated cells such as neuronal-like and astrocyte-like tumor cells. While our study does not ascribe function to these cells, we have observed hybrid-incompletely differentiated cells in glioblastoma patients capable of regulating neuronal activity within the tumor microenvironment[50]. Furthermore, we have previously demonstrated in glioma, that differentiated cells can contribute to tumor progression by releasing paracrine factors that maintain glioma stem cell identity and regulate tumor progression[51]. Our findings may also provide an explanation for the highly chemo-resistant nature of EPN, where persistent and non-/slowly-dividing malignant cells resist conventional chemotherapies, and failure to eliminate cycling progenitor-like clones may facilitates EPN regrowth.

## EXTENDED DATA FIGURE LEGENDS

**Extended Data Figure 1. E14.5 forebrain developmental patterns for 10xmultiome (snRNA + snATAC) data. (a)** Differential gene expression between ZFTA-RELA fusion and WT in radial glia (RGC) using bulk RNA-Seq. The marked genes are the 93 genes closely associated with ZR oncogenic activity. **(b)** Validation of our cell type characterization using public single-cell RNA datasets (Bella et al.,Nature 2021) and examining cell cycle signaling. **(c)**Identification of key gene signals in the forebrain using spatial ATAC-Seq data (Zhang et al., Nature 2023). The yellow region indicates open chromatin. **(d)** Investigation into whether ZFTA-RELA directly binds to PLAGL1/2 motifs using cognate site identification (CSI). **(e)** Identification of the top 5 known cell type markers per cluster, as reported in previous papers (Table S2). **(f)** Examination of gene expression patterns (snRNA) of the PLAG family in the E14.5 forebrain. **(g)**Analysis via box plot of Plagl1 motif (MA1615.1) activity between radio glial cells (RGC) and neuronal cells (Neuron). **(h)**Display of a scatter plot illustrating the average motif activity signal for each motif between RGC and Neuron. **(i)**Examination of peak signal in the target genes between RGC and Neuron. Small orange vertical lines indicate ZFTA-RELA binding sites, collected from previous Chip-seq studies.

**Extended Data Figure 2. In the mouse IUE model, encompassing ZFTA-RELA and GBM tumors, patterns were analyzed using 10xmultiome (snRNA + snATAC). (a)** Copy number alterations (CNAs) were examined in snRNA from mouse IUE ZFTA-RELA samples (IUE-ZR, n=3). Red indicates copy number gain, while blue signifies loss. **(b)** CNA patterns in snATAC were assessed in IUE-ZR. **(c)** Each sample’s proportion of malignant and non-malignant cells was determined, along with the distribution of malignant cells in UMAP. **(d)** Identification of the top 5 known cell type markers per cluster in IUE-ZR. **(e)** Proportions of potential cell types identified for IUE-ZR multiple groups. **(f)** CNA patterns in snRNA were assessed in mouse IUE GBM sample. **(g)** Identification of the top 5 known cell type markers per cluster in IUE GBM. **(h-j)** Gene expression patterns of the PLAG family (Plag1, Plagl1, and Plagl2) in IUE-ZR. **(k)** The chromatin accessibility ratio of ZFTA-RELA fusion binding sites between RGC-Like and Neuronal-Like cells was investigated. **(l)** A box plot illustrates Plagl1 motif (MA1615.1) activity between RGC-Like and Neuronal-Like cells. **(m)** A scatter plot shows the average motif activity signal for each motif between RGC-Like and Neuronal-Like cells.

**Extended Data Figure 3. Define malignant cells in Human EPN multiome data (n=19, PF=10, ZR=6, EWP=3). (a)** Copy number alterations (CNAs) in snATAC profiles of human EPN samples (n=19). The red color signifies a gain in copy numbers, while the blue color indicates a loss in copy numbers. **(b)** The proportion of malignant and non-malignant cells in each sample. **(c)** Estimation of target genes of the PLAGL family (PLAGL1 and PLAGL2) and comparison of the number of target genes between EWP and ZR group EPNs. This schematic diagram illustrates a concept for predicting target genes of the PLAGL family. For instance, if a PLAGL family motif is mapped to the promoter region of a gene, that gene is identified as a target gene of the PLAGL family. The left Venn diagram represents the number in the promoter region, while the right one illustrates the number of target genes. **(d)** The heat map displays the peak signal intensity of the PLAGL family motif within the promoter region between EWP and ZR groups. **(e)** Results of gene ontology (GO) enrichment analysis in biological processes (BP) using total PLAGL family target genes between EWP and ZR groups. The green color represents shared pathways, while the orange color denotes ZR-specific pathways.

**Extended Data Figure 4. Human EPN cell type annotation. (a)** Identification of the top 5 known cell type markers per cluster from malignant cells in ST-EPN. **(b)** Identification of the top 5 known cell type markers per cluster from malignant cells in PF-EPN. **(c-d)** Potential cell types were identified for ST-EPN and PF-EPN per sample. The x-axis represents the sample names. **(e)** Showing the top 5 known cell type markers for each cluster in ST-EPN ZFTA-RELA (ZR) samples, data obtained by extracting ZR subgroups from ST-EPN integrated malignant cells and re-clustering the data. **(f)** Bar plot showing cell type proportion per cell type. **(g)** Comparing the cell type annotations of the ZR group data by our lab with the annotations from another lab. **(h)** PLAG family gene expression patterns in ZR group. **(i)** PLAGL family motif activity signal in ZR group. **(j)** Cell cycle alteration signals in malignant cells of ZR group. Cluster 5 (C5) showed high cell cycling signals. **(k)** Validating our cell type characterization using public single-cell RNA datasets (Trevino et al.,Nature 2021) and examining cell cycle signaling.

**Extended Data Figure 5. Mouse IUE ZR EPN tracker-seq data and cell type annotation. (a, d, g)** Cell type annotation of Tracker-seq data where cells contain single or multiple lineage barcodes. Tracker-Seq P4 is a low tumor purity sample, Tracker-Seq #1 and #2 are high tumor purity samples. **(c)** Integrated the technically replicated Tracker-seq data for Tracker #2 sample. **(b, e, h)** Signature of ZR within the cells. **(f, i)** The cellular pattern indicative of the cell cycle signature.

## EXTENDED DATA TABLES

**Table S1 | Dataset overview.**

**Table S2 | Developmental brain cell type markers.**

**Table S3 | Cell type, Plagl motif, pseudotime Inference, and cell cycling study for E14.5 forebrain development.**

**Table S4 | Comprehensive analysis of mouse IUE using multiome-seq data from ZR EPN and GBM tumors.**

**Table S5 | Comparative differential analysis of human ependymoma subgroups using multiome data.**

**Table S6 | Cell type annotation for malignant cells and ZR signal.**

**Table S7 | Tracker-seq analysis overview for moue IUE ZR EPN.**

**Table S8 | Pseudotime and cell-cell communication for ZR EPN tumors.**

## METHODS

### In Utero Electroporation

All animal procedures in this study were performed with St. Jude Institutional Animal Care and Use Committee approval. IUE was performed as described previously, and plasmids were prepped with the NucleoBond® Xtra Maxi Plus EF kit (Takara Biosciences). After anesthesia with 4.5% isoflurane, pregnant mice at E16.5 were subjected to abdominal incision to expose the uterus. DNA plasmid cocktail (1ug/ul pBCAG-HA-ZR^FUS1^, 1ug/ul pbCAG-eGFP-Luciferase, 1.5ug/ul pX330-sg*Tp53*, 2ug/ul GLAST-PBase, 1.5ug/ul m*Plagl1* sgRNA, 0.5 ug/ul TrackerSeq library, FastGreen dye) was injected into the lateral ventricles with a glass pipette. Electric pulses were then delivered to the embryos by gently clasping their heads with forceps-shaped electrodes. Six 33 V pulses of 55 ms were applied at 100-ms intervals. The uterus was then repositioned into the abdominal cavity, the abdominal wall was sutured, and the skin was stapled. Following birth, pups were monitored for clinical signs of tumor growth (seizures, circling, head doming, etc) as well as by MRI every 2 weeks. At endpoint, mice were collected for nuclei isolation or IF staining. Mice for nuclei isolation were perfused with 10mL cold PBS and tumors were frozen in isopentane and stored at –80C.

### Radial Glial Cell Isolation

Radial glia cells were previously isolated from *Ink4a* knock out mice with GFP expressed from the *Blbp* promoter using the Worthington Papain Dissociation system (LK003150). Cells were grown in neural basal medium (Invitrogen) supplemented with sodium pyruvate, glutamine, B27, N2, bFGF (10 ng/mL), and rhEGF (20 ng/mL). Cells were grown on treated cell culture dishes coated with Matrigel (Corning). RGCs were made ZFTA-RELA positive via lentivirus generated by the Viral Vector Core at St. Jude. Mouse tumor cells were seeded into 10-cm plates 24 hours prior to infection and were infected with lentivirus with 8 μg/mL polybrene for 24 hours. Infected cells were selected with 2 μg/mL puromycin for 3 days. ZFTA-RELA expression was confirmed by western blot. SgRNAs for *Plagl1* were generated by the Center for Advanced Genome Engineering with AddGene 52961 backbone, which included RFP. ZFTA-RELA positive RGCs were infected by the same method and were sorted for RFP via a BD FACSAria Fusion system. Over 90% knock out was confirmed by targeted deep sequencing.

### RNA-seq and ATAC-seq Analysis

RNA-seq and ATAC-seq analysis were performed using Genialis Expression software (https://www.genialis.com) deployed locally on St Jude HPC infrastructure. Briefly, the RNA-seq pipeline run on the Genialis platform comprised the following steps. The raw reads were filtered to remove adapters and poor-quality reads using BBDuk (v37.9; https://sourceforge.net/projects/bbmap/). The resulting reads were mapped to the reference genomes (ENSEMBL 92) using STAR (v2.7.0; RRID:SCR_015899). FeatureCounts (v1.6.3; RRID:SCR_012919) was used for gene expression level quantification followed by DEseq2 (RRID:SCR_000154) for differential gene expression analysis. Low-expressed genes with expression count summed over all samples below 10 were filtered out from the input matrix to DESeq2. The paired-end reads from ATAC-seq were trimmed using BBDuk (v37.9) and mapped to the reference genome mm10 using Bowtie2 (v2.3.4.1). MACS2 (v2.1.1.20160309) was then used to call peaks on the aligned reads using *P* value cutoff of 0.01 (parameters –shift -75 –extsize 150 –nomodel –call-summits –nolambda –keep-dup all -*P* = 0.01)

### TrackerSeq library generation and validation

TrackerSeq library cloning was carried out generally as described in Bandler et al[34]., Briefly, the pCAG-SacB plasmid was digested with BstXI and the 8bit barcode was cloned into it using NEBuilder HiFi master mix (6 reactions total), followed by isopropanol purification. Purified reactions were electroporated into Endura DUOs (Lucigen) using a MicroPulser with program Ec1 (Bio-Rad). Four electroporations were carried out, then recovered for 1 hr at 37C in 2ml of recovery media. Next, cells were plated overnight at 32C on 245mm plates (Corning). The following morning, plates were scraped and harvested using LB, and library plasmids purified using Endofree midiprep kits (Qiagen). For validation, 10ng of the library plasmid prep was amplified using 2xPhusion (NEB) and sequenced by the Hartwell Center for Genome Sequencing Facility at St. Jude.

### Cognate Site Identification DNA binding assay

HEK293T cells were transiently transfected with HA-tagged ZFTA-RELA fusion plasmids using lipofectamine 2000 reagent according to the protocol (Thermo Fisher 11668019). Following expression, cells were lysed with RIPA buffer (Thermo Fisher 89900) and spun down to collect supernatant. A DNA library (Integrated DNA Technologies) containing randomized central regions of 20 bp flanked by constant sequences complimentary to primers was converted to dsDNA and brought to 74ng/μL before being combined 1% w/v BSA, 500ng/μL poly dI-DC (Thermo Fisher 20148E), 1% NP-40 (Thermo Fisher 85124), and 10X PBS. ZFTA-RELA positive and negative cell-lysates were incubated with this mixture for 1 hour at room temperature. The mixture was then added to anti-HA beads (Thermo Fisher 88836) washed in binding buffer (10X PBS, 1% BSA, 1% NP-40,) and incubated for 30 mins. Solutions were washed in binding buffer and aspirated on a magnetic plate three times before being resuspended in a PCR master mix (Lucigen Econo Taq 2X 30035-1 and custom primers). Library fragments attached to beads were amplified on a Bio-Rad thermal cycler and then purified using the New England Biolabs Monarch PCR & DNA clean-up kit (T1030L). Eluted DNA library fragments from each sample were diluted to a concentration of 74ng/μL and checked on a gel before being incubated with cell lysates again. Following three rounds of this incubation and amplification, all purified library fragments from each round for each sample were given a unique barcode and a sequencing adapter and then sequenced on a NovaSeq short-read sequencing amplicon kit yielding ∼500 million reads. Sequencing results were sorted by barcode and the 20bp library regions selected in each sample were ranked by enrichment and normalized to fusion negative lysate samples. Primer Sequences: forward 5’-CTGATCCTACCATCCGTGCT-3’, reverse5’-CCGCTCGGTACGAAGCTG-3’

### Nuclei isolation

10-30mg of tissue was cut from human and IUE tumors and was input into the 10x Nuclei Isolation Kit (PN-1000494). Kit instructions were followed but lysis buffer incubation time was increased to 15 minutes to isolate quality nuclei. Mouse embryonic forebrain was isolated with the 10x Demonstrated Protocol CG000366 RevD, but lysis buffer incubation was decreased to 2 minutes. Nuclei were resuspended in nuclei buffer, counted via hemocytometer, and 10,000 were loaded onto the 10x Chromium Chip. Libraries were assessed via Agilent TapeStation and sent for sequencing at St. Jude’s Hartwell Center on a NovaSeq 6000. Gene Expression libraries had 400 million reads with a 28-10-10-90 cycle configuration and ATAC libraries had 500 million reads with a 50-8-24-49 cycle configuration.

### Single cell multiome data processing

For human scMultiome, raw snRNA-seq and snATAC-seq reads were aligned to the GRCh38 (10x Genomics, refdata-cellranger-arc-GRCh38-2020-A-2.0.0) reference genome and produced ‘filtered_feature_bc_matrix’ and fragment files using ‘cellranger-arc count’ (10x Genomics, v.2.0.1, https://www.10xgenomics.com). For mouse scMultiome, the raw sequence reads were aligned to the mm10 (10x Genomics, refdata-cellranger-arc-mm10-2020-A-2.0.0) reference genome. After running ‘cellranger-arc’, the ‘filtered_feature_bc_matrix’ and fragment files produced, and the files containing profiles of both snRNA- and snATAC-seq data were read into R with the Seurat (v.4.3.0, https://satijalab.org/seurat) function Read10X. To obtain the final list of barcodes, we retained the cells that passed the quality control filters in both the snRNA- and snATAC-seq assays using Signac (v.1.9.0, https://github.com/timoast/signac) multiome pipeline. Next, we performed separate normalization and integration processes for snRNA- and snATAC-seq data (Detailed individual processing is shown below). Last, we computed the weighted nearest neighbour (WNN) graph with the FindMultiModalNeighbors function using both integrated data modalities. We used dimension reduction 1:30 integrated snRNA-seq and 2:30 integrated snATAC-seq for this analysis. And we performed non-linear dimensionality reduction of the resulting WNN graph using the RunUMAP function of Seurat. Finally, we obtained clusters with the FindClusters function using the WNN graph, setting the resolution = 0.4.

### snRNA processing of multiome

The gene expression count data extracted from multiome gene UMI count matrix and was further processed using the Seurat (v.4.3.0, https://satijalab.org/seurat). We first filtered features detected in a minimum of three cells. Next, we selected high-quality cells based on below cutoffs such as more than 500 genes, cells with more than 2000 unique molecular identifiers counts (nCount_RNA), cells with less than 10% of mitochondrial genes, more than 0.7 of log10GenesPerUMI that equal to log10(nFeature_RNA)/log10(nCount_RNA), and less than customized maximum quantile for nFeature_RNA (< as.integer(Q[2]+1.5*IQR); Q=quantile(probs = c(.2, .8)), IQR=Q[2]-Q[1]) and nCount_RNA (< as.integer(Q[2]+1.5*IQR); Q=quantile(probs = c(.1, .9))). We performed doublet estimation using the DoubletFinder (v.2.0.3, https://github.com/chris-mcginnis-ucsf/DoubletFinder), but did not find clear evidence of cell doublets biasing our unsupervised analysis and therefore did not apply doublet filtering. Thereafter, expression matrices of high-quality cells were normalized using SCTransform v2 method. Principal component analysis (PCA) was performed based on the 3,000 most variable features (SelectIntegrationFeatures). To integration all of samples, we merged all single snRNA data, performed PCA for the merged data, and batch correction using Harmony (v.0.1.1, https://github.com/immunogenomics/harmony) with default parameters with Seurat SCTransform. We used the first 30 Harmony embeddings for UMAP visualizations and clustering analysis. To partition cells into clusters, we constructed a shared-nearest neighbour graph based on Harmony embeddings via the ‘FindNeighbors’ and ‘FindClusters’ functions in Seurat (dims = 30, resolution = 0.3). Cluster-specific marker genes were identified by comparing cells of each cluster to cells from all other clusters using ‘FindAllMarkers’ function in Seurat with Wilcoxon rank sum test. Differentially expressed genes were defined based on fold change, gene expression in percentage of cells and adjusted p-value cut-offs.

### snATAC processing of multiome

The UMI count matrix and fragment data, which were generated by ‘cellranger-atac count’, was further processed using the Signac (v.1.9.0, https://github.com/timoast/signac). We filtered out cells with less than 3,000 or more than as.integer(Q[2]+1.5*IQR) sequencing fragments, and discarded cells with a TSS enrichment less than 4 or more than 15, nucleosome signal less than 4, and blacklist fraction less than 4. Next, peak calling was then performed on the MACS2 tool (v.2.2.6, https://github.com/macs3-project/MACS/tree/v2.2.6) through the CallPeaks function of the Signac package. And remove peaks on nonstandard chromosomes and in genomic blacklist regions. To integration all snATAC-seq data, we first created a unified set of peaks to quantify in each dataset using GenomicRanges package, and filtered peaks less than 5 or larger than 10,000 in peak widths. And the fragment of each sample was recalled based on the unified peaks. And gene activity and motif activity were calculated based on the new peak and fragment data per sample. Next, we merged all snATAC data and performed TF-IDF normalization and LSI-dimensionality reduction using the RunSVD function from the Signac package. Non-linear dimensionality reduction was performed using the RunUMAP function with 2:50 LSI components. Last, we processed integration and batch correction using the Harmony (v.0.1.1), generated non-linear dimensionality using the RunUMAP function, and performed clusters using via the ‘FindNeighbors’ and ‘FindClusters’ functions in Seurat.

### Estimation of motif activity from snATAC-seq data

Motif/transcription factor chromatin accessibilities (motif activity) was computed for set of 841 TFs (combined mouse and human TFs) from the JASPAR 2022 using the RunChromVAR function in Signac (v.1.9.0) and differential motif activity was computed with the FindMarkers function. Motif enrichment analysis was performed on the differentially accessible regions with the FindMotif function.

### Annotation of cell types for multiome

Cell-type annotations were assigned to WNN UMAP partitions and initial clusters in multiome using manual collection of marker gene set (**Table S2**) and the ScType annotation tool (v.2022-01-12, https://github.com/IanevskiAleksandr/sc-type). Since we used same cells that have snRNA and snATAC in multiome, the cell type definition was based on snRNA gene expression with marker gene set and choose the top five known marker gene per cell type to present in the figures.

### Tracker-Seq data processing for mouse IUE

Raw Tracker-Seq data was processed using cellranger (v.7.2.0, https://www.10xgenomics.com) with mm10 (10x Genomics, refdata-cellranger-mm10-2020-A-2.0.0) reference genome. In the calling counts, we set --force_cell = 5000 to increase medium gene counts per cell. After that, filtered_feature_bc_matrix data was used to filter pool cells and normalized in Seurat. We selected high-quality cells based on below cutoffs such as more than 800 genes, cells with more than 2000 unique molecular identifiers counts (nCount_RNA), cells with less than 20% of mitochondrial genes, more than 0.8 of log10GenesPerUMI that equal to log10(nFeature_RNA)/log10(nCount_RNA), and less than customized maximum quantile for nFeature_RNA (< quantile(probs = 0.998) and no more than 8,000) and nCount_RNA (< quantile(probs = 0.998)). Next, we performed doublet estimation using the DoubletFinder (v.2.0.2) and kept ‘Singlet’ cells. After that, normalized by SCTransform v2 method with regress out mitochondrial genes in Seurat (v.4.3.0). Lastly, high quality cells were used to run PCA, UMP, and clusters in Seurat. Cell type annotation was performed using ScType (v.2022-01-12) with manually collected marker gene sets (**Table S2**).

### Isolate high-confidence lineage barcodes of matching cells from Tracker-seq

Processing of TrackerSeq barcode reads, we followed Bandler et al., 2021[34] methods (https://github.com/mayer-lab/Bandler-et-al_lineage). First, trimming the sequences to the left and right of the lineage barcodes and discarding shorter than 37 bp lineage barcodes from FASTQ file. Next, cell barcodes were extracted from the corresponding Seurat object of the dataset to establish a cell barcode whitelist. These extracted cell barcodes and UMIs were then appended to the read names of the lineage barcode FASTQ files. Subsequently, the resulting FASTQ files were processed to generate a sparse matrix in CSV format. Within this matrix, rows represented cells identified by individual cell barcodes, while columns represented lineage barcodes. To reduce false positive lineage barcodes, we only kept the lineage barcodes with conserved nucleotide ‘…CTG…ACT…GAC…TGA…CTG…ACT…GAC…’, and without any ‘N’ in the lineage barcodes. For multiple replicate sequencing of a sample, we used lineage barcodes that are shared with all replicate data. In downstream analyses, we only used cells with a single lineage barcode in a single cell and filtered out cells with multiple lineage barcodes in a single cell.

### Classification of malignant cells

To classify cells as malignant or non-malignant, we inferred genome-wide copy-number alterations (CNAs) using InferCNV (v.1.14.2, https://github.com/broadinstitute/infercnv/blob/master/inst/NEWS) with the default parameters. InferCNV was run at the sample level with after integrated counts matrix of snRNA- and snATAC-seq data, separately. The reference used immune cells and microglial cells. We first define the CNAs as <= 0.8 (loss) or >= 1.2 (gain) based on InferCNV value and filtered CNA regions (hidden Markov model outputs) that < 20 number of genes for detecting large region chromosomal CNAs. Next, we calculate CNA ratio per CNA region. Last, we estimated malignant cells based on large CNA region (>= 100 CNA genes in 5 Mb region) or CNA ratio (> 0.6) per sample.

### ZFTA-RELA fusion signal signatures

Single cell ZFTA-RELA (ZR) fusion signature was calculated based on 93 ZR driver genes[17, 18, 39] using the AddModuleScore function in Seurat (v.4.3.0). Based on in-house benchmark test, a score > 0.2 indicates confidence, 0.1-0.2 indicates uncertainty, and < 0.1 indicates distrust. Negative controls were tested using normal samples, ependymomas without ZR fusion and glioblastoma.

### Cell cycling and non-cycling signatures

Cell cycle score was calculated using the CellCycleScoring function in Seurat (v.4.3.0). Based on the function, the cell cycle spited three different phase such as G1, G2M, and S. To define cell cycle and non-cycling, we re-calculated cell cycle score based on cell cycle markers inner Seurat (cc.genes.updated.2019) using the AddModuleScore function in Seurat. And named the re-calculated score as cc.score. Based on internal testing, we set scores >0.2 in cc.score or S.score or G2M.score as cell cycling cells, and <0.1 as non-cycling cells.

### Motif target genes and gene enrichment

We utilized the Signac functions ‘keepStandardChromosomes’ and ‘promoters’ to identify promoter regions in protein-coding genes. Overlapping peaks were merged using the default parameters of the ‘findOverlaps’ function in the R package. We then employed Signac to extract peaks associated with the target motif and retained only those peaks that had at least 5 reads per cell and presented in at least 100 cells. Finally, the cleaned peaks were annotated using ChIPseeker (v1.34.1, https://github.com/YuLab-SMU/ChIPseeker), and only protein-coding genes for peaks corresponding to promoter regions were considered. Gene enrichment analysis was performed using the enrichGO function from clusterProfiler (v4.6.0, https://github.com/YuLab-SMU/clusterProfiler). For the EPN subgroups, we first extracted the target subgroup cells and then performed the same steps as above.

### Construction of pseudo-time trajectories

Monocle (v.2.30.0, https://cole-trapnell-lab.github.io/monocle-release) was used to convert SCT-normalized sn/scRNA-seq data from Seurat object into a cell dataset object (CDS). Next, differentially expressed genes were identified using M3Drop package (v1.28.0, https://github.com/tallulandrews/M3Drop), and dimensionality reduction was performed using the ‘reduceDimension’ function of Monocle (parameter: norm_method=’none’, reduction_method=’DDRTree’). Finally, pseudo-time ordering was carried out by setting the root cell type. And ‘plot_cell_tracjectory’ function of Monocle was used to visualize pseudo time landscape for single cells.

### Progenitor and lineage score of malignant cells

The progenitor and lineage score were calculated following the method of Filbin et al., 2018[30]. We only used malignant cells that defined by InferCNV (descripted in the methods) and set cell cycling enrichment cell type as progenitor (nIPC-/CycProg-Like) to start in the top. Progenitor score was estimated as expression of the progenitor shared program minus the maximal expression of the two differentiation programs, and the differentiated cells were further classified based on average gene expression differences to differentiate between astrocyte-like and neuron-like lineages.

## DATA AND CODE AVAILABILITY

Data generated for this study will be available through the Gene Expression Omnibus (GEO, accession no. GSE269937). All other data are available in the manuscript or the supplementary figures or supplementary tables. All data needed to evaluate the conclusions in the paper are present in the paper and/or the extended data.

## ACKNOWLEDGMENTS

This study was supported by a NCI Cancer Center Support Grant, P30 CA021765, St. Jude Children’s Research Hospital Research Collaborative on Transcription Regulation in Pediatric Cancer Grant, Alex’s Lemonade Stand Foundation ‘A’ Award. This work is supported by R01NS128184, R01CA280203, R01CA284455, U01CA281823, DOD-IDEA (CA220510), and an DOD-IMPACT (CA220247) award (to SCM). SCM is supported by funding from the National Brain Tumor Society and CERN Foundation. This work was also supported by US National Institutes of Health grants R35-NS132230, R01-NS124093, R01-CA223388 to B.D.; the National Cancer Institute Cancer Target Discovery and Development grant U01-CA217842 to B.D. The authors would like to acknowledge Jackie Norrie and the Single Cell Core for her training and allowing us to use their Chromium machine. We would also like to acknowledge the Christian Mayer lab for their generous donation of Tracker-Seq plasmids. We are grateful to Dr. Aseem Ansari lab for helping us with application and analysis of the CSI technology to study ZR. We are also grateful to Dr. Carol Schuurmans for her generous donation of Plagl related plasmids and valuable advice. Several other St. Jude core facilities were also instrumental in this work, including the Hartwell Sequencing Center, the Viral Vector core, and the Flow Sorting core.

## AUTHOR CONTRIBUTIONS

A.K., H.S., K.B., B.D., and SCM., were involved in the conception and design of the project. A.K., S.I., H.C., B.H., and N.L., and performed experiments. A.K., S.I. and E.E. were involved in mouse care. H.S. performed single cell sequencing analysis and visualization, prepared data tables, and managed single cell data. H.S. and S.C.M. organized figures. S.V. performed bulk sequencing analysis and visualization. N.L. performed CSI experiment and visualization. K.L. performed flow sorting for TrackerSeq experiments. P.C., S.M.P., J.P.C., and M.W. expanded and validated the TrackerSeq library. S.C.M., A.K., H.S., N.L., P.C., and S.V. wrote the manuscript.

## COMPETING INTERESTS STATEMENT

The authors have no conflicts of interest to disclose

## MATERIALS AND CORRESPONDENCE

This study did not generate new reagents. The data generated within this study are available within the article, the supplementary data files, and at the Gene Expression Omnibus accession number GSE269937. Correspondence and requests for materials should be addressed to corresponding author Stephen C. Mack, Benjamin Deneen, and Kelsey C. Bertrand.

## Notes

The authors of this manuscript have no conflicts of interest to disclose.

### Competing Interest Statement

The authors have declared no competing interest.

