## Extended Data Figures for "Dominant Malignant Clones Leverage Lineage Restricted Epigenomic Programs to Drive Ependymoma Development"

### Extended Data Figure 1

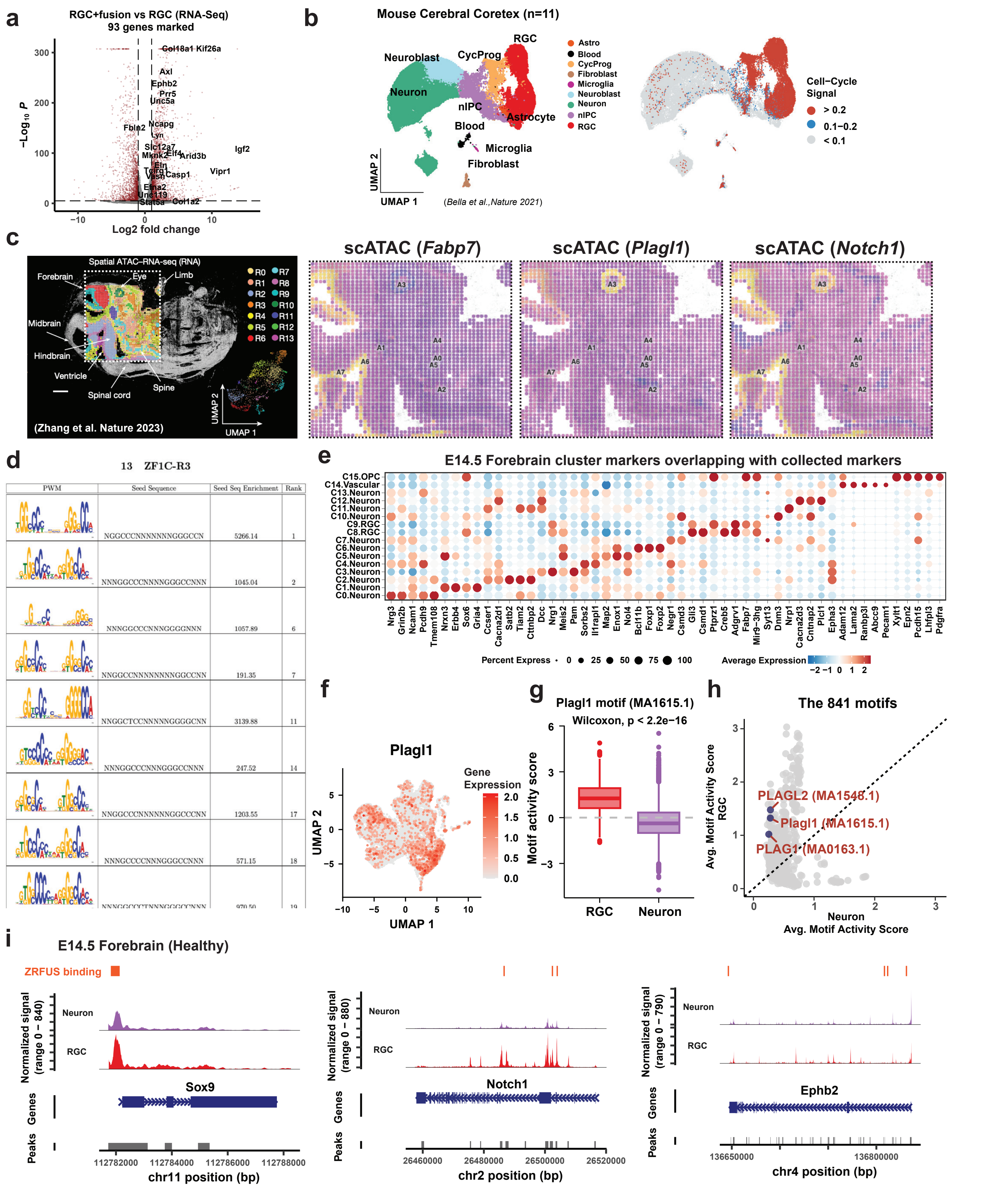


**Extended Data Figure 1. E14.5 forebrain developmental patterns for scMultiome (snRNA + snATAC) data. (a)** Differential gene expression between ZFTA-RELA (ZR) fusion and WT in radial glia (RGC) using bulk RNA-Seq. The marked genes are the 93 genes closely associated with ZR fusion oncogenic activity. **(b)** Validation of our cell type characterization using public single-cell RNA datasets (Bella et al.,Nature 2021) and examining cell cycle signaling. **(c)** Identification of key gene signals in the forebrain using spatial ATAC-Seq data (Zhang et al., Nature 2023). The yellow region indicates open chromatin. **(d)** Investigation into whether ZR directly binds to PLAGL1/2 motifs using cognate site identification (CSI). **(e)** Identification of the top 5 known cell type markers per cluster, as reported in previous papers (Table S2). **(f)** Examination of gene expression patterns (snRNA) of the PLAG family in the E14.5 forebrain. **(g)** Analysis via box plot of Plagl1 motif (MA1615.1) activity between radio glial cells (RGC) and neuronal cells (Neuron). **(h)** Display of a scatter plot illustrating the average motif activity signal for each motif between RGC and Neuron. **(i)** Examination of peak signal in the target genes between RGC and Neuron. Small orange vertical lines indicate ZR binding sites, collected from previous Chip-seq studies.

### Extended Data Figure 2

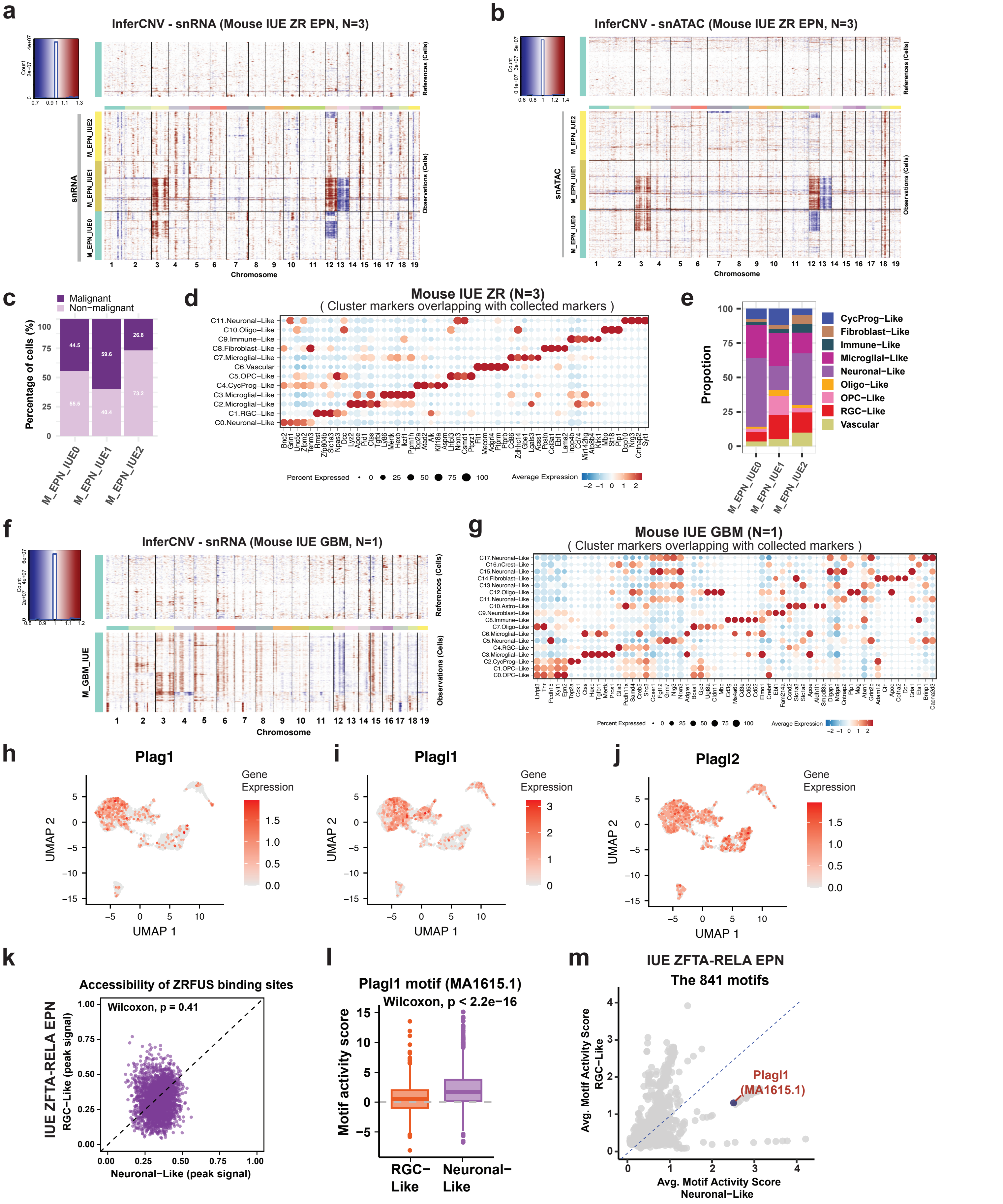


**Extended Data Figure 2. scMultiome landscape of ZFTA-RELA and GBM tumors in mouse IUE model. (a)** Copy number alterations (CNAs) were examined in snRNA from mouse IUE ZR samples (n=3). Red indicates copy number gain, while blue signifies loss. **(b)** CNA patterns in snATAC were assessed in IUE ZR. **(c)** Each sample's proportion of malignant and non-malignant cells was determined, along with the distribution of malignant cells in UMAP. **(d)** Identification of the top 5 known cell type markers per cluster in IUE ZR. **(e)** Proportions of potential cell types identified for IUE ZR multiple groups. **(f)** CNA patterns in snRNA were assessed in mouse IUE GBM sample. **(g)** Identification of the top 5 known cell type markers per cluster in IUE GBM. **(h-j)** Gene expression patterns of the PLAG family (Plag1, Plagl1, and Plagl2) in IUE ZR. **(k)** The chromatin accessibility ratio of ZR fusion binding sites between RGC-Like and Neuronal-Like cells was investigated. **(l)** A box plot illustrates Plagl1 motif (MA1615.1) activity between RGC-Like and Neuronal-Like cells. **(m)** A scatter plot shows the average motif activity signal for each motif between RGC-Like and Neuronal-Like cells.

#### Extended Data Figure 3

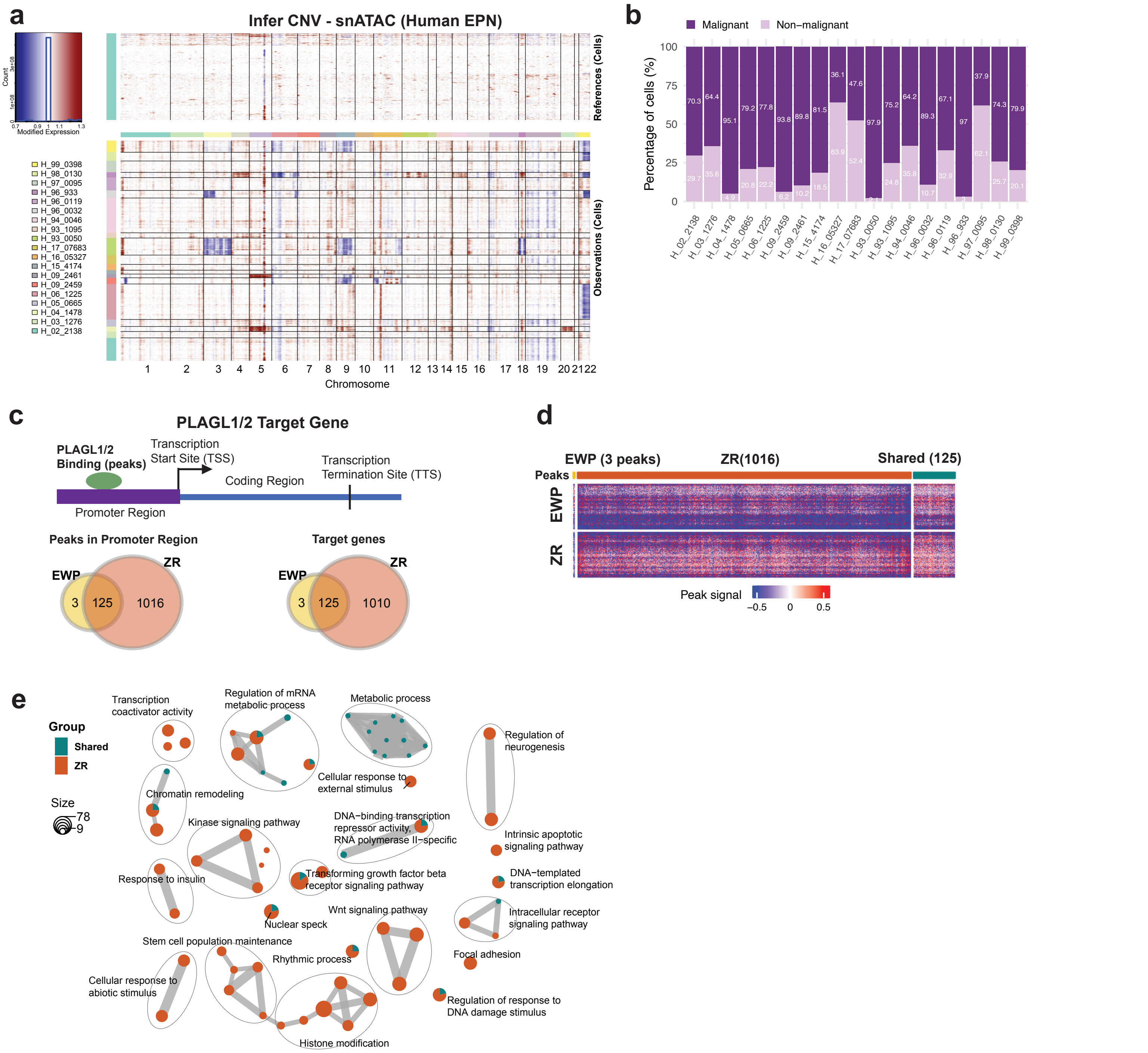


**Extended Data Figure 3. Define malignant cells in Human EPN scMultiome data (n=19, PF=10, ZR=6, EWP=3). (a)** Copy number alterations (CNAs) in snATAC profiles of human EPN samples (n=19). The red color signifies a gain in copy numbers, while the blue color indicates a loss in copy numbers. **(b)** The proportion of malignant and non-malignant cells in each sample. **(c)** Estimation of target genes of the PLAGL family (PLAGL1 and PLAGL2) and comparison of the number of target genes between EWP and ZR group EPNs. This schematic diagram illustrates a concept for predicting target genes of the PLAGL family. For instance, if a PLAGL family motif is mapped to the promoter region of a gene, that gene is identified as a target gene of the PLAGL family. The left Venn diagram represents the number in the promoter region, while the right one illustrates the number of target genes. **(d)** The heat map displays the peak signal intensity of the PLAGL family motif within the promoter region between EWP and ZR groups. **(e)** Results of gene ontology (GO) enrichment analysis in biological processes (BP) using total PLAGL family target genes between EWP and ZR groups. The green color represents shared pathways, while the orange color denotes ZR-specific pathways.

### Extended Data Figure 4

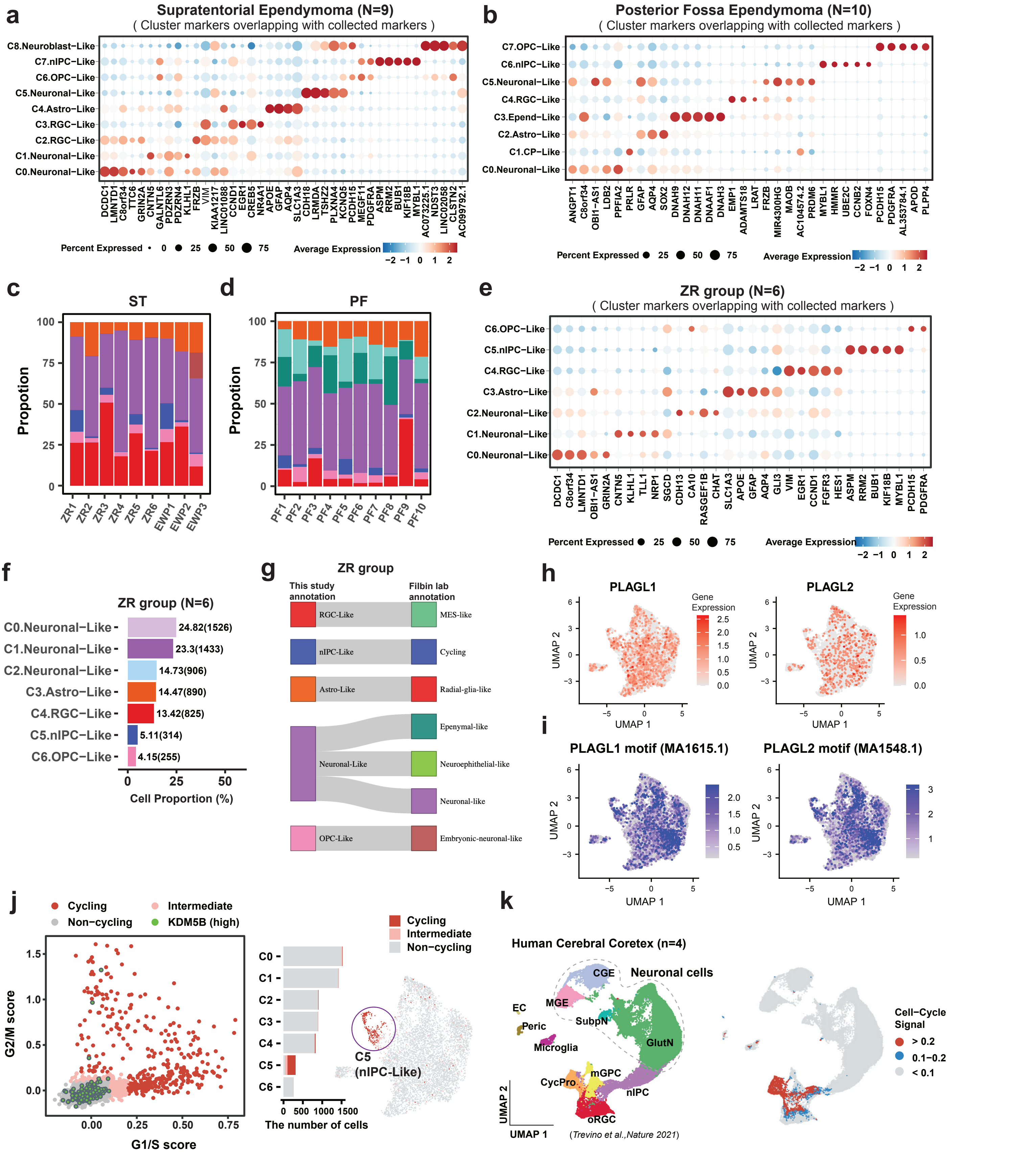


**Extended Data Figure 4. Human EPN cell type annotation. (a)** Identification of the top 5 known cell type markers per cluster from malignant cells in ST-EPN. **(b)** Identification of the top 5 known cell type markers per cluster from malignant cells in PF-EPN. **(c-d)** Potential cell types were identified for ST-EPN and PF-EPN per sample. The x-axis represents the sample names. **(e)** Showing the top 5 known cell type markers for each cluster in ST-EPN ZR samples, data obtained by extracting ZR subgroups from ST-EPN integrated malignant cells and re-clustering the data. **(f)** Bar plot showing cell type proportion per cell type. **(g)** Comparing the cell type annotations of the ZR group data by our lab with the annotations from another lab. **(h)** PLAG family gene expression patterns in ZR group. **(i)** PLAGL family motif activity signal in ZR group. **(j)** Cell cycle alteration signals in malignant cells of ZR group. Cluster 5 (C5) showed high cell cycling signals. **(k)** Validating our cell type characterization using public single-cell RNA datasets (Trevino et al.,Nature 2021) and examining cell cycle signaling.

### Extended Data Figure 5

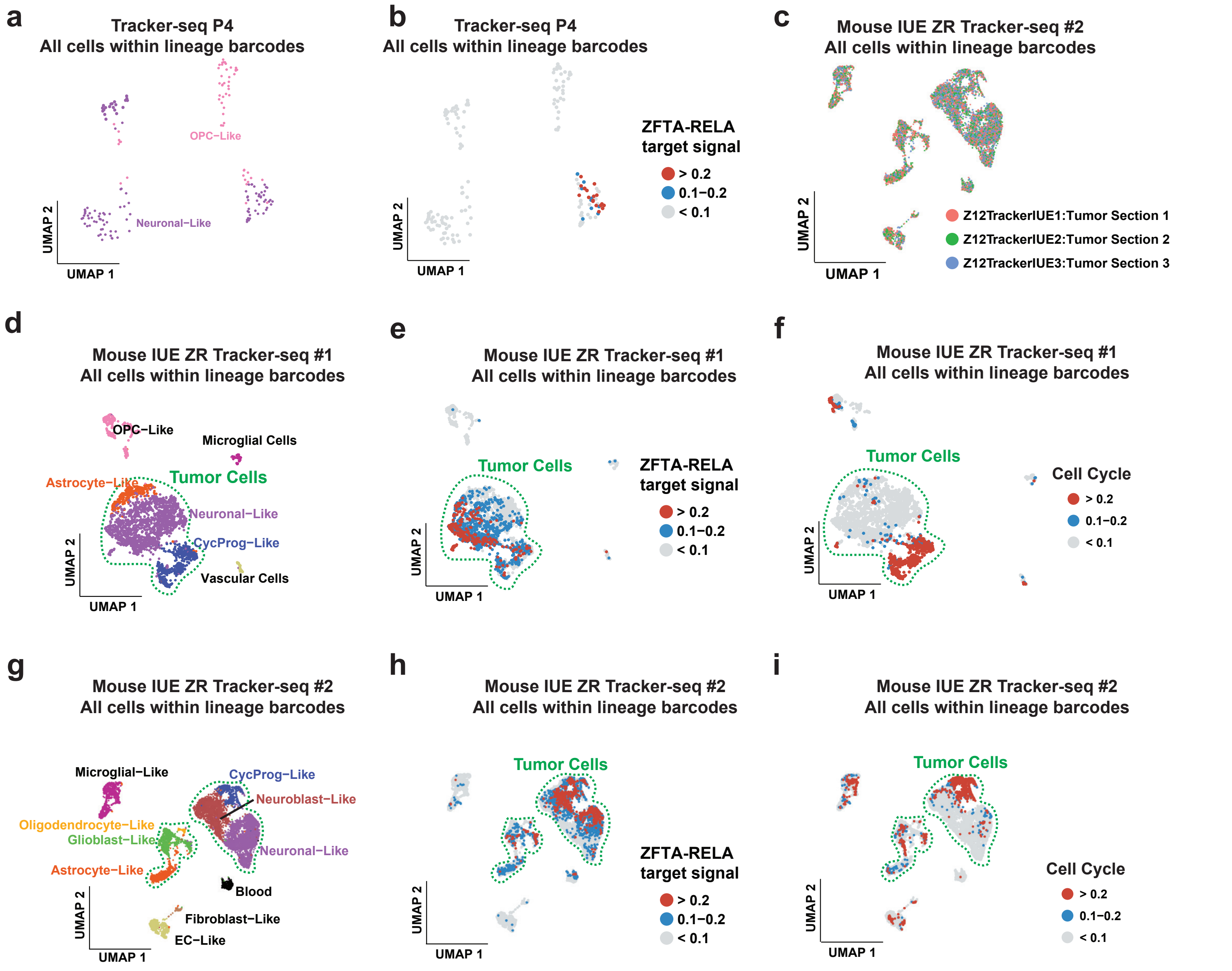


**Extended Data Figure 5. Mouse IUE ZR EPN tracker-seq data and cell type annotation. (a, d, g)** Cell type annotation of Tracker-seq data where cells contain single or multiple lineage barcodes. Tracker-Seq P4 is a low tumor purity sample, Tracker-Seq #1 and #2 are high tumor purity samples. **(c)** Integrated the technically replicated Tracker-seq data for Tracker #2 sample. **(b, e, h)** Signature of ZR within the cells. **(f, i)** The cellular pattern indicative of the cell cycle signature.
